# Ezh2 restrains macrophage inflammatory responses, and is critical for neutrophil migration in response to pulmonary infection

**DOI:** 10.1101/2020.07.24.219154

**Authors:** Gareth B. Kitchen, Thomas Hopwood, Thanuja G. Ramamoorthy, Polly Downton, Nicola Begley, Tracy Hussell, David H. Dockrell, Julie E. Gibbs, David W. Ray, Andrew S.I. Loudon

## Abstract

Mucosal immunity is critical to survival, with huge attention at present due to the Coronovirus pandemic. Epigenetic factors are increasingly recognized as important determinants of immune responses, and EZH2 closest to application due to the availability of highly-specific and efficacious antagonists. However, very little is known about the role of EZH2 in the myeloid lineage, with some conflicting reports. Here we show EZH2 acts in macrophages to limit inflammatory responses to activation, and selective genetic deletion results in a remarkable gain in protection from infection with the prevalent lung pathogen, pneumococcus. In contrast, EZH2 is required for neutrophil chemotaxis, and animals lacking neutrophil EZH2 show increased susceptibility to pneumococcus. In summary, EZH2 shows complex, and divergent roles in different myeloid cells, likely contributing to the earlier conflicting reports. Compounds targeting EZH2 are likely to impair mucosal immunity, however, may prove useful for conditions driven by pulmonary neutrophil influx, such as adult respiratory distress syndrome (ARDS).

**Digest:** Epigenetic control of mucosal immunity is important, and has translational relevance with the advent of inhibitor drugs now in the clinic for cancer indications. Here we show divergent role for EZH2 in macrophages and neutrophils. Loss of EZH2 in macrophages results in a gain of inflammatory and immune function, and protection from pneumonia. However, EZH2 is required for neutrophil chemotaxis, resulting in impaired anti-bacterial defence. We show that inhibition, or loss of EZH2 in macrophages results in a gain of immune function, with increased responses to infectious mimics such as LPS. However, the impact was far more dramatic in-vivo, with striking protection from the consequences of infection with pneumococcal bacteria. Loss of EZH2 resulted in a gain in activity of a number of inflammatory signaling cascades, including NFkB, PPARg, and IRFs1, and 7. This widespread macrophage re-programming varied between macrophages sites of origin, with the greatest impact seen in peritoneal macrophages which resulted in emergence of a new population of MerTK low cells. In contrast, in the neutrophils loss of EZH2 greatly impairs motility, and chemotaxis. This results in dramatic impairment of immune responses to the same pneumococcal infection. Extension of these studies to the mucosal epithelium revealed that EZH2 in bronchoalveolar epithelial cells had no impact on responses to infection with influenza. Taken together EZH2 plays diverse roles in the myeloid lineage, with profound impacts on inflammatory responses. The most striking observation was the difference seen between macrophages and neutrophils. EZH2 inhibition is likely to greatly impair mucosal immunity.

**Impact Statement:** Here we show a striking, but highly cell-type specific impact of the EZH2 methyltransferase on inflammatory, and anti-infective circuits; inhibition of EZH2 in macrophages augments macrophage cytokine production, but by impairing neutrophil migration impairs anti-bacterial responses.

## Introduction

Macrophages are widely distributed through vertebrate tissues and play essential roles in homeostasis, and in response to injury. Therefore, it is unsurprising that these cells have been implicated in the pathogenesis or host response to a wide-range of prevalent human diseases, including fibrosis, osteoporosis, obesity, type 2 diabetes and atherosclerosis (1-3). Activation of macrophages is critical for an appropriate response against an invading pathogen. Alveolar macrophages, for example, are located in the alveolar spaces of the airways and provide the first line of defense against respiratory pathogens, such as the gram-positive bacteria *Streptococcus pneumoniae*. Recognition of bacteria, and many other pathogens, is facilitated by toll-like receptor (TLR) pathways. TLRs are a group of cell-surface and endosomally-located pattern-recognition receptors (PRRs), which, in conjunction with adaptor proteins, activate signaling cascades to induce a cellular response. Multiple TLRs, through MyD88-dependent, and independent cascades converge to activate NFkB, AP-1 and IFN regulatory factors (IRFs).

Epigenetics, the study of chromatin modifications which alter DNA availability, has been known for many years to be crucial in the formation of a functional immune response (4). Recently, attention has focused on the role of methylation of histone3Lys27 (H3K27), mediated by the polycomb-2 (PRC2) complex, comprising Suz12, Eed and Ezh2, with EZH2 as the catalytic subunit. PRC2 is a powerful determinant of differentiated cell state, and by silencing gene expression, limits trans-differentiation. EZH2 has been intensely studied in cancer (5, 6), and has gained attention in generating a functional, adaptive immune response (7, 8). Macrophage switching between functional states has been identified to be dependent on reversible trimethylation of histone H3K27 (9).

Several recent studies have employed genetic targeting of EZH2 or pharmacological EZH2-selective inhibitors to demonstrate an important role in the regulation of inflammatory responses. These investigations have yielded contradictory results. Pro-inflammatory outcomes were documented following suppression of EZH2 in inflammatory bowel disease models (10-12) and Kras-driven lung cancer inflammation (13). In contrast, Zhang et al demonstrated that EZH2 in the macrophage drives experimental autoimmune diseases including experimental autoimmune encephalomyelitis (EAE) and colitis. In this latter study, loss of EZH2 reduced inflammatory responses to stimulus and so reduced disease severity, pointing the way to pharmacological inhibition of EZH2 as a therapeutic strategy in multiple sclerosis (14). These discrepancies may result from differential impacts on different cell types, and gene expression programs involved in immune response elaboration, and require further analysis.

Here we show that the actions of EZH2 in the myeloid lineage is complex, with a role inhibiting the pro-inflammatory responses of macrophages, and enabling efficient neutrophil migratory capacity. This dichotomy of action may also explain the apparent contradictory outcomes of earlier studies of the role of EZH2 in inflammation. The complex actions of EZH2 on innate immune responses are important to consider with progression of drugs targeting this powerful lineage maintenance, epigenetic regulator into the clinic for cancer indications, and suggestions of a role in managing autoimmune disease such as multiple sclerosis.

## Results

### Deletion of EZH2 increases inflammatory cytokine production in multiple macrophage lineages – Fig 1

We assessed the efficacy of genetic targeting, using cultured bone-derived macrophages (BDMs) from myeloid-targeted mice and showed loss of H3K27 trimethylation (Fig 1A). We also showed loss of EZH2 gene expression in alveolar, peritoneal, and bone marrow derived macrophages (FIG S1). We also employed an EZH2 inhibitor in wild-type BMDMs (UNC1999), and showed resulting loss of H3K27me3, similar to the genetically targeted cells (Fig 1A).

**Figure 1:**
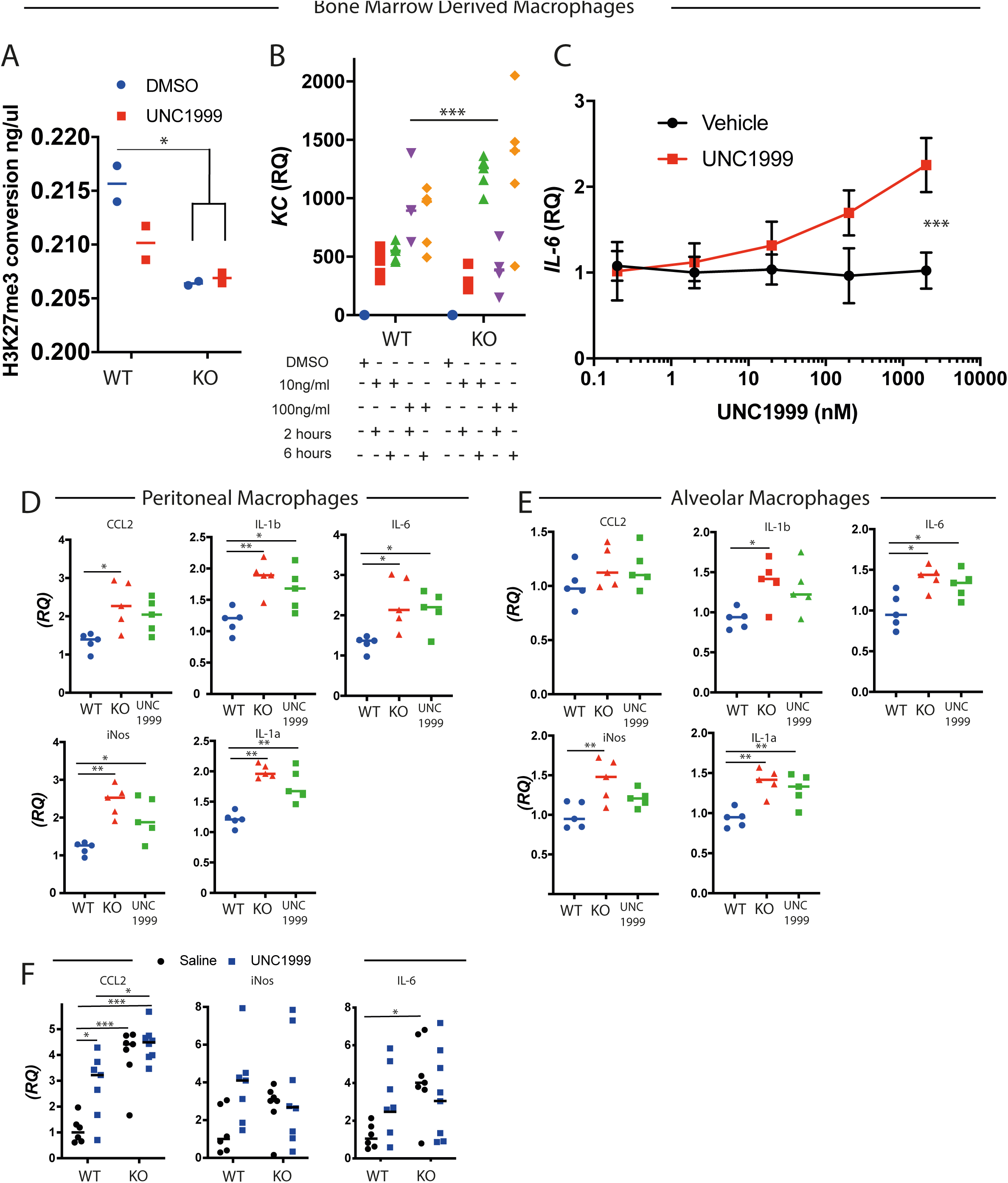
All experiments performed using LysMcre driver. WT = EZH2 fl/fl KO = LysM x EZH2 fl/fl. A) Functional assay of EZH2 in nuclear lysates from BMDMs derived from Ezh2^fl/fl^ LysM-cre mice treated for 48h with either DMSO or 2μM UNC1999. Methyltransferase assay confirmed that genetic targeting of EZH2 in macrophages reduced the conversion of H3K27me1 to H3K27me3, as indicated by loss of trimethylation in Bone Marrow Derived Macrophages (BMDMs) from these mice. This effect was mimicked by use of the EZH2 inhibitor UNC1999 in WT cells, but was not additive to the effect of loss of tri-methylation in Ezh2^fl/fl^ LysM-cre derived cells (n=4, 2-way ANOVA, post-hoc Tukey’s multi-comparison test *p=<0.05, median plotted). B) Isolated macrophages from bone marrow and treated with different doses of LPS for 2 or 6 hours shows that these cells are highly sensitive to time and concentration of treatment. Ex-vivo BMDMs from EZH2^-/-^ macrophages and littermate controls were treated with LPS for 2 or 6 hours at a dose of 10 or 100ng/ml and the cytokine response assessed by RNA quantification. (n= 5, median plotted, 2-way ANOVA, post-hoc Tukey’s multi-comparison test. *** p<0.001). C) UNC1999 causes a dose dependent increase in IL6 expression in BMDMs Ex-vivo BMDM cells were incubated with either UNC1999 or vehicle at the indicated concentrations for 48 hours before stimulation with LPS (100ng/ml) for 2 hours. quantification of IL6 mRNA by qRT-PCR. (n= 3, 2-way ANOVA, post-hoc Tukey’s multi-comparison test. *** p<0.001). D) Peritoneal exudate cells (PECs) and E) Alveolar macrophages were harvested from EZH2^fl/fl^ (WT) or EZH2^fl/fl^.LysMcre (KO) mice and treated with vehicle for 1 hour, before exposure to 10ng/ml LPS for 6h. Additionally, cells from EZH2^fl/fl^ mice were pre-treated with 2μM UNC1999 for 1h prior to LPS (labelled UNC1999). Cytokine response was measured by qRT-PCR for a panel of inflammatory cytokines. These data show elevated cytokine responses in EZH2-disrupted PECs, an effect mimicked by treatment with UNC1999, but no significant effect on responses of alveolar macrophages. Ex-vivo analysis of PECs and alveolar macs revealed small increase in inflammatory mediator expression in PECs and UNC1999 showed a similar effect to the genetic loss of EZH2 in the cells, with minimal change in alveolar cells (n= 5, median plotted, 2-way ANOVA, post-hoc Tukey’s multi-comparison test. P< 0.05 **p<0.01, *** p<0.001). F) In-vivo responses were assessed in EZH2^fl/fl^ (WT) or EZH2^fl/fl^. LysMcre (KO) mice which were either pre-treated with either vehicle, or 2μM UNC1999 by an intra-peritoneal injection (IP) of 50mg/kg per day for 2 days before a single IP injection of LPS 1mg/kg. Peritoneal cells were washed out after 2 hours, and inflammatory mediators quantified by qRT-PCR. mRNA was quantified relative to GAPDH, and was normalised to vehicle treated control mice given LPS. The results show an enhanced inflammation in EZH2-disrupted cells, with no additive effect of the drug to that of the genetic model. (n=6-8, * P,0.05 vs vehicle treated mice, 2-way ANOVA, post-hoc Tukey’s multi-comparison test.).

Next, we generated mice lacking EZH2 in the macrophage lineage, for in-vivo infection studies, by using the C3CXR-cre driver line. We tested the in-vitro response of Ezh2-deficient BMDMs to 10 or 100ng/ml LPS for 2 or 6h (Fig1B, and Fig S2). The pattern of gene expression revealed a complex interaction between genotype LPS concentration, and duration of exposure, but in general cytokine mRNA species were greater at 6 than 2 hours post activation, and responses to LPS were greater in the absence of macrophage EZH2 (Fig 1B). In particular the responses of KC were striking, with duration of LPS exposure exquisitely impacting the LPS induction in the absence of EZH2. This finding is similar to a previous report, but the impact of EZH2 is more complex than a simple gain or loss of global inflammatory response (14). We then moved on to test the EZH2 inhibitor UNC1999 (15) on wild-type BMDMs, which showed a dose-dependent increase in expression of *IL-6* in BDMs post LPS-treatment (Fig 1C). We extended these analyses to two further EZH2 inhibitors and found in both cases dose-dependent induction of LPS-induced *IL6* (Fig S3).

Macrophages are found in multiple tissue compartments and exhibit a variety of different phenotypes (16). To check the extent of the wider actions of EZH2 we isolated peritoneal exudate cells (PECs) (Fig 1D) and alveolar macrophage (AMs) (Fig 1E) from EZH2^f/f^ (WT) and LysM-cre.EZH2^f/f^ (KO) mice. The isolated cells were treated with 10ng/ml LPS for 6h, before analysis of a panel of LPS-induced inflammatory genes. Here we observed that either genetic loss of EZH2, or pharmacological inhibition amplified the cytokines responses to macrophage activation.

Because the actions of EZH2 on the responses of isolated macrophages from different sites showed some differences, and some of the responses we saw conflicted with earlier publications (14) we opted to move to in-vivo analyses. Here, we treated Ezh2^fl/fl^ LysM-cre and Ezh2^fl/fl^ mice with either UNC1999 or vehicle for two days before intraperitoneal LPS (1mg/kg) (Fig 1F). Genetic loss of EZH2 resulted in a target gene selective gain in response to LPS injection, with CCl2, and IL-6 showed significantly increased responses. The UNC1999 significantly increased CCL2 induction, but only in the EZH2f^/f^ control mice, and only a marginal, and non-significant impact on responses of *iNOS* and *IL6*.

### EZH2 deletion enhances RelA activation in response to TLR-4 activation – Fig 2

Studies on pre-defined panels of inflammatory genes may miss wider impacts of the epigenetic changes driven by EZH2. Accordingly, to investigate the wider impact of *Ezh2* deletion on expression of immune related genes in BMDMs we used a pre-made CodeSet (Mouse inflammation panel V2 Nanostring™ (Fig 2A; Fig S4). We saw that the major changes in gene expression was between the EZH2f/f controls, and the LysM-cre.EZH2 cells, with LPS exposure of the cells exerting a dominant effect of the gene expression profile (Fig 2A). As many of the differentially regulated genes under basal conditions were components of the TLR-NFkB system (Supp Table1), we draw on some examples (Fig 2B). The TLR signaling system employs a number of overlapping elements (Fig 2C), and so to refine the parts of the cascade functionally affected by EZH2 we used a TransAm assay (Invitrogen). Here we were able to show that ligands for TLR4 caused increased activation of both NFkB, and IRF3, but ligands specific for TLR3 (PolyI:C), or TLR2 (Pam3csk4) were not impacted by loss of EZH2 (Fig 2D).

**Figure 2.**
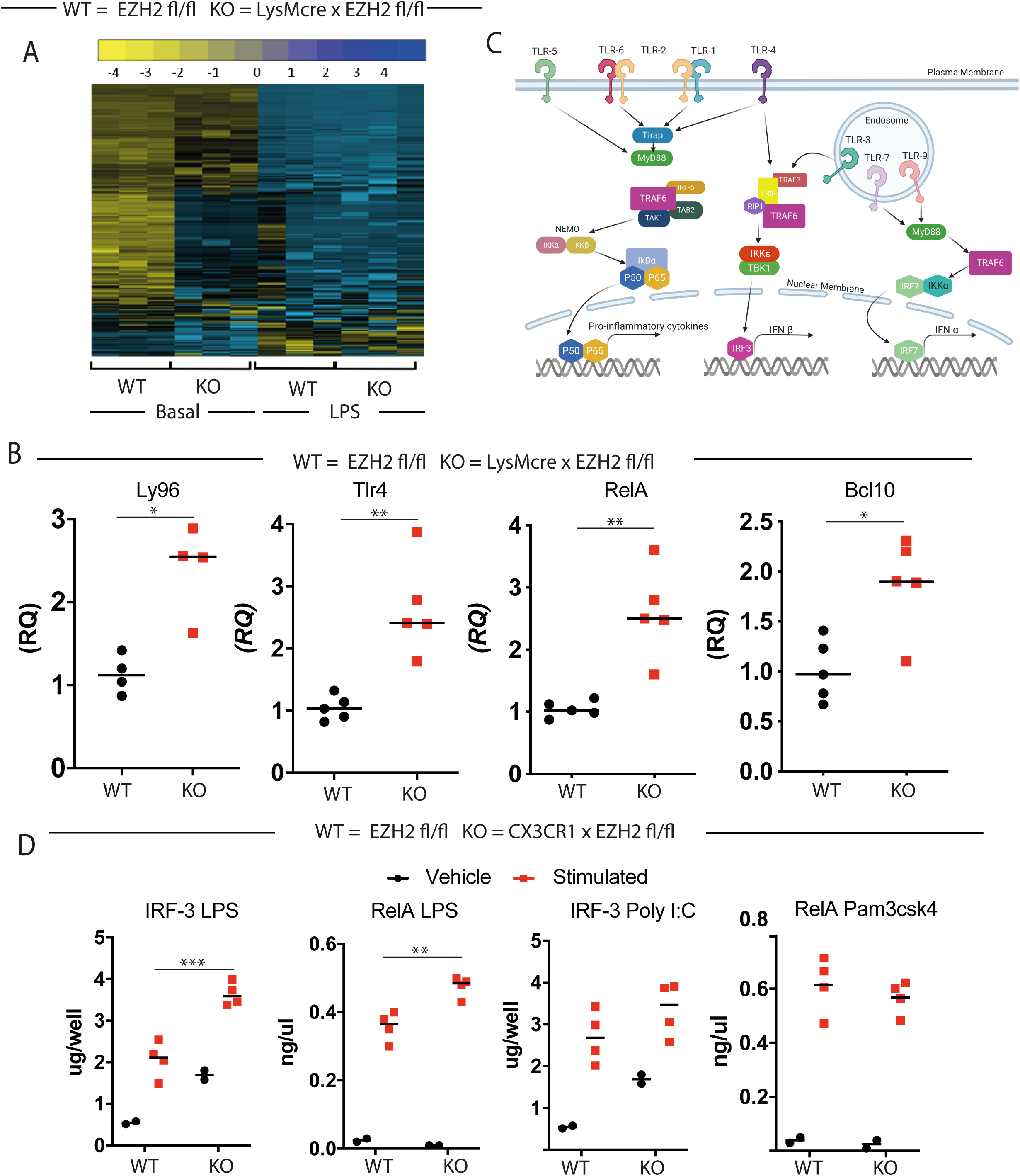
Selective impact of EZH2 deletion on TLR4 signaling. A) To investigate the consequences of EZH2 deletion on immune pathways a Nanostring analysis was performed comparing BMDMs from EZH2f/f.LysMcre-cre (KO) mice with EZH2^f/f^ (WT) littermate controls. This analysis was undertaken using the Mouse Inflammation Panel V2 Nanostring code-set for inflammatory gene expression. Cells were differentiated in-vitro, and analysed under un-stimulated basal conditions. The RCC files (NanoString compressed data filetype) were then analyzed in the nSolver™ software package (NanoString Technologies). The heat-map shows the three replicates. B) This revealed increased transcript levels of Ly96, TLR4, RelA and BCL10, all components of the TLR4 Pathway analysis suggested increased expression of TLR4 pathway components, and accordingly the specific transcripts identified were plotted. (n=4-5, median plotted, Mann-Whitney *U*-test *P< 0.05 **p<0.01, ***p<0.001) C) Diagram of the main TLR signaling pathways, highlighting overlap between TLR1/2, TLR3, and TLR4 cascades. D) To investigate the potential impacts on other TLR pathways we used a TLR agonist panel to challenge each TLR individually in the absence of EZH2. Cells were incubated with the TLR1/2 ligand Pam3csk4, the TLR3 ligand PolyI:C, or the TLR4 ligand LPS for 2 hours before harvest and analysis using the TransAm system. Transam Assays revealed that in the absence of EZH2 an LPS challenge leads to increased activity of both RelA (P65) and IRF-3, but that PolyI::C and Pam3csk4 did not result in altered transcription factor activation. (n= 2-4, median plotted, 2-way ANOVA, post-hoc Tukey’s multi-comparison test. P< 0.05 **p<0.01, *** p<0.001)

To extend our analyses further we used RNA-sequencing to profile responses in BMDMs lacking EZH2 (LysM-cre.EZH2^f/f^ compared to EZH2^f/f^). PCA plots revealed tight clustering and strong separation by both genotype and LPS treatment (Fig 3A). Again, we observed a greater separation under basal conditions than after LPS activation; a similar result to that seen in the Nanostring analysis above (Figure 2 A,B). Volcano plots visualize genotype differences under basal (Fig 3B) and LPS-stimulated conditions (Fig3C). These show that the major down-regulated transcript is Xist, a non-coding RNA necessary for X-chromosome inactivation. Rather more genes were increased than decreased in expression by deletion of EZH2, as expected from loss of an inhibitory epigenetic enzyme. A number of cytokines, chemokines, and NFkB signaling components were increased by loss of EZH2 under the LPS-activated state (Fig 3C).

**Figure 3.**
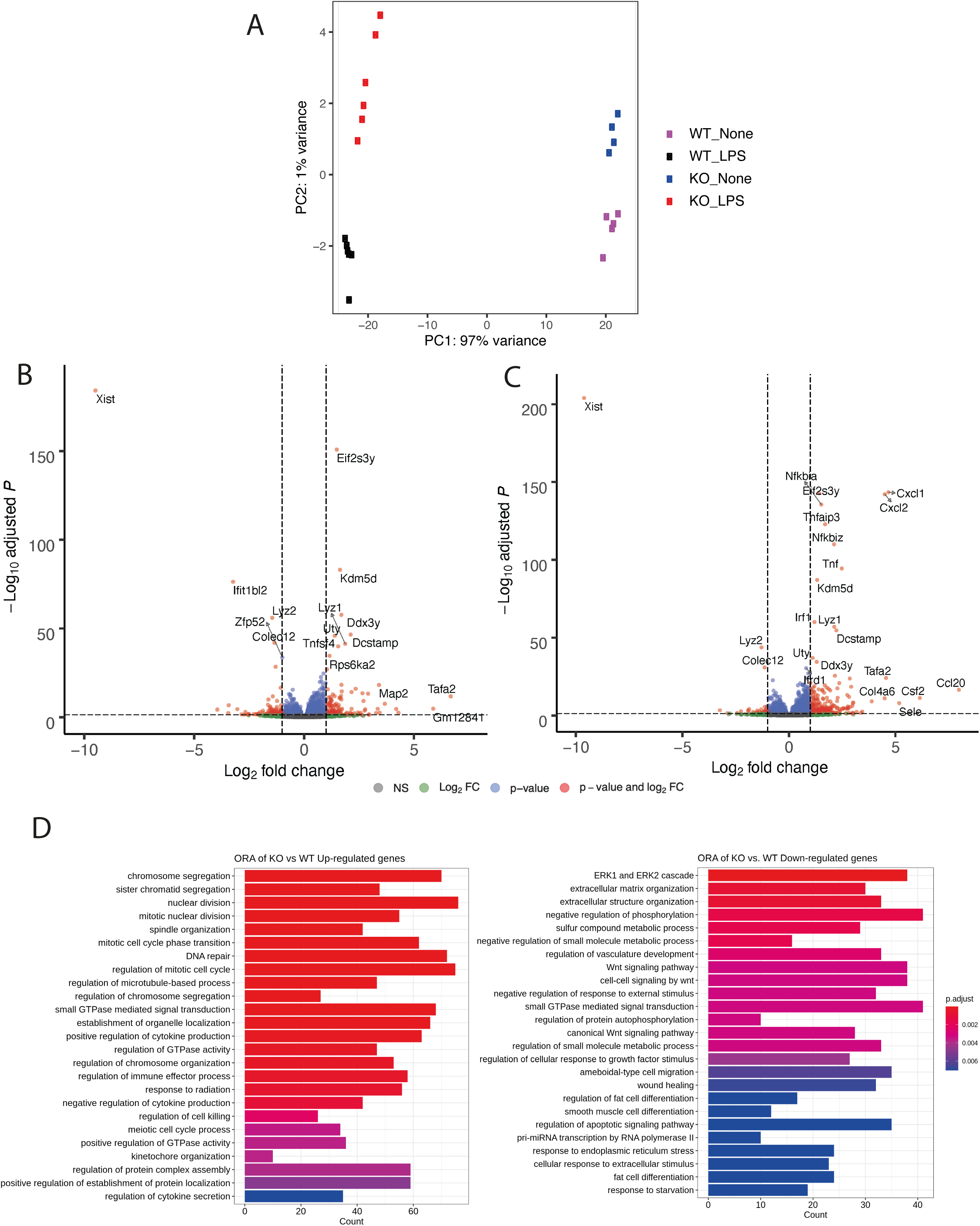
RNA-SEQ analysis of BMDMs lacking EZH2. BMDMs from EZH2^fl/fl^ (WT) or EZH2^fl/fl^.CX3CR1-cre (KO) mice were differentiated in vitro, and then treated with either vehicle or LPS for 2 hours prior to harvest and RNA-SEQ. A) Principal components analysis revealed clear differentiation by genotype under basal, and activated conditions. B,C) Volcano plots comparing BDMs derived from EZH2^fl/fl^ (WT) or EZH2^fl/fl^.CX3CR1-cre (KO) mice in the basal state and following LPS administration. 55,385 genes measured, Log2 fold change cutoff 1, p-value cutoff 0.05. Only a minority of the quantified 55,385 genes showed differential expression by genotype, with similar numbers of genes showing up or down-regulation. F) Gene ontology over-representation analysis (ORA) identified a number of processes that were regulated by EZH2 under basal conditions in the macrophage. The colour blue to red indicated increasing statistical significance and count is expressed as a histogram on the x-axis. Further analysis in Fig S5A-E.

To analyse these differences further we used a gene ontology approach with gene ontology over-representation analysis (ORA)(17). We selected gene lists of EZH2-differentially regulated genes under basal conditions, as we saw a greater effect in the absence of LPS activation. Gene ontology terms that were enriched in the upregulated gene lists included a surprising cluster associated with cytoskeletal function, chromosome segregation, and motility. In addition, we observed enrichment of terms involved in cytokine and chemokine regulation. In contrast the gene ontology terms associated with the down-regulated gene list were involved in signaling, and extracellular matrix regulation (Fig 3D).

From the EZH2 regulated gene lists we performed further analysis using Ingenuity Pathway Analysis (IPA: Qiagen) (Fig S5A-E). We saw similar upstream regulators in both genotypes with 19 of 20 of the most significant genes being the same.

We saw that multiple upstream signaling cascades were regulated by loss of EZH2, with no single dominant pathway (Fig S5A,B). In basal comparison (KO vs WT, no LPS), the only transcriptional regulators detected as enriched upstream regulators with expression log ratio > 0.5 were PPARg (up), and Tcf7, Id3 and meis1 (down) (Fig S5C). In the LPS-activated samples several transcriptional regulators showed significant enrichment and differential expression dependent on EZH2. We observed that Irf7, Irf1, Spib, PPARa, Nfkbia and Nfkbiz were expressed at higher levels in the KO compared to WT (Fig S5D), this result is similar to that revealed by the nanostring analysis in Fig 2 and fig 3 B and C. volcano plots. This finding provides a useful extension to the NFkB functional analyses (Fig 2D), and further supports dysregulation of NFkB circuits by loss of EZH2. Expression analysis is largely consistent with upregulation of these regulators, particularly IRF7 and IRF1 (Fig S5E).

### Targeted deletion of EZH2 in pulmonary macrophages provides protection against bacterial infection

Analyses to date had revealed a complex network of EZH2 effects in myeloid cells, and so to determine the physiological significance we moved to further in-vivo testing. To evaluate the in-vivo macrophage-specific responses, we focused on use of the macrophage-specific Cx3CR1-cre driver to target EZH2 deletion to macrophage lineages (Fig4A)(18). Loss of Ezh2 in macrophages differentially affects surface marker expression in cells derived from different compartments (Fig 4A), with the emergence of a distinct population of MerTK low macrophages in the peritoneum that were not seen in the alveolar macrophages. In differentiation studies on bone marrow we also did not observe the emergence of a low MerTK population of macrophages. MerTK is an important cell-surface receptor on macrophages, tending to drive activation of pro-inflammatory genes, and so the low MerTK population likely is an anti-inflammatory population. However, as they were only seen in the peritoneal compartment, we did not pursue analysis further.

**Figure 4.**
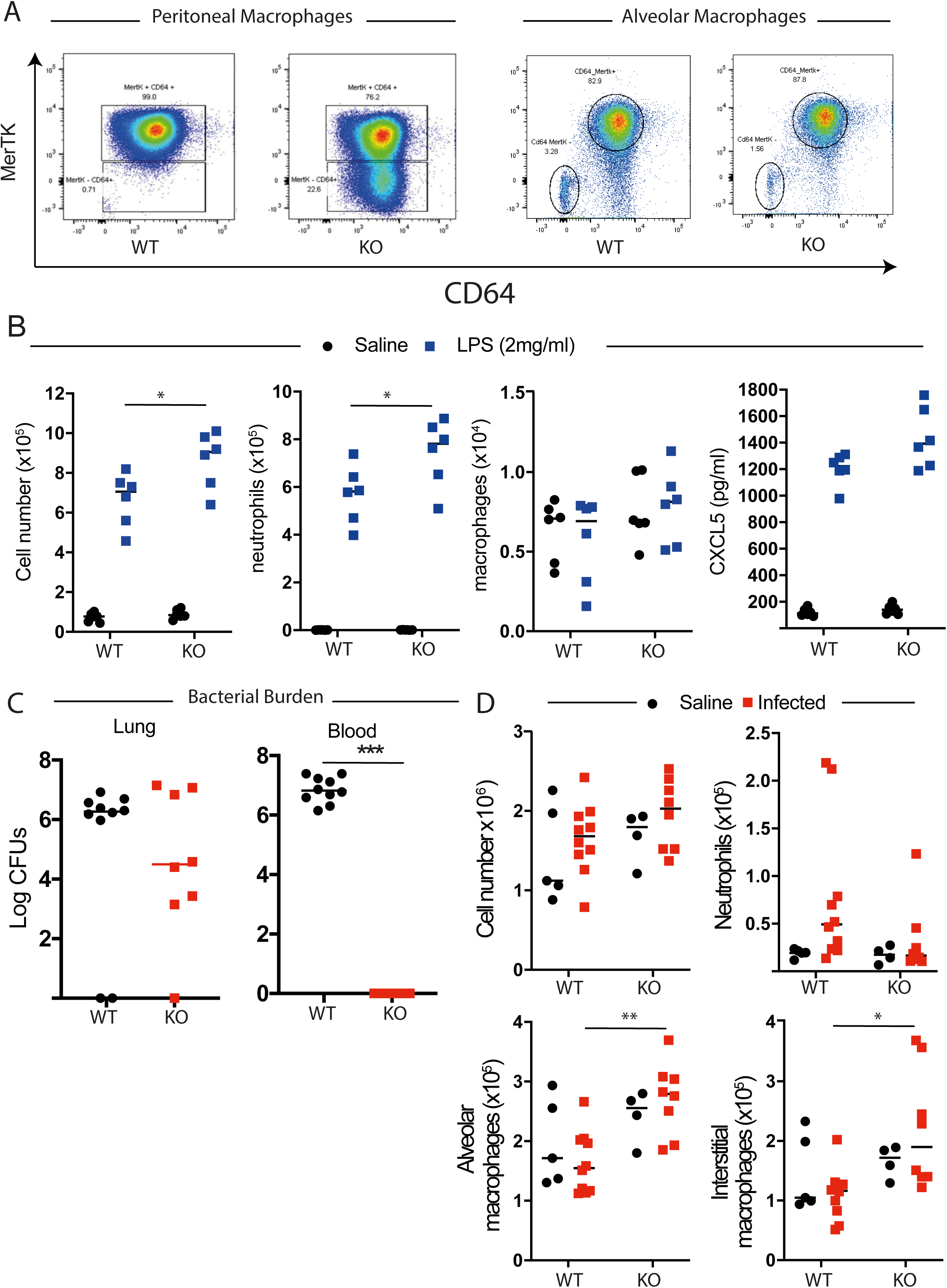
Inflammatory and immune function of EZH2 in macrophages. A To investigate in-vivo consequences of enhanced inflammatory activation we investigated LPS activated macrophages from two compartments, the peritoneal cavity and alveolar spaces. Macrophage-specific targeting of EZH2 was undertaken using CX3CR1-cre EZH2^fl/fl^ mice. FACS analysis showed that the peritoneal macrophages displayed an extra ‘population’ of macrophages without EZH2 that lack the surface marker MerTK. This is a phenotype purely limited to the peritoneum and is not present in the alveolar spaces which could not be distinguished by genotype on cell surface markers. B Pulmonary in-vivo inflammatory responses in CX3CR1-cre EZH2^fl/fl^ mice. Mice were subject to nebulized LPS, and after 4 hours killed. The bronchoalveolar lavage (BAL) was analysed by flow cytometry, and the immune cell fractions are plotted. This showed limited genotype-dependent differences in the number of cells infiltrating into the lung after challenge with slight changes in neutrophil numbers, (n=6, median plotted, 2-way ANOVA, post-hoc Tukey’s multi-comparison test. P< 0.05 **p<0.01, *** p<0.001) C Pneumococcal responses in EZH2^fl/fl^ or EZH2^fl/fl^.CX3CR1-cre mice were compared. All animals were inoculated with pneumococcus, and then analysed 48 hours later. Bacterial counts (colony-forming units, CFUs) in the lung, and peripheral blood were measured, and are plotted. Following streptococcal challenge to the lung there were limited differences in the CFUs present in lung homogenate, but in contrast bacterial levels in the blood clearly shows a very strong phenotype, with no detectable CFUs in the blood stream at the time of cull from EZH2^fl/fl^.CX3CR1-cre mice. (n=8-10, median plotted, Mann-Whitney *U*-test *P< 0.05 **p<0.01, ***p<0.001). D Myeloid responses to inflammatory activation in EZH2^fl/fl^.CX3CR1 in macrophages. Mice of the indicated genotypes were subject to nebulized LPS as before, and then killed after 24 hours. The lungs were homogenized, and myeloid cell content measured by FACS. Alveolar and interstitial macrophages were distinguished by cell surface markers (Fig S7). These results show that EZH2^fl/fl^.CX3CR1 (KO) mice can rapidly re-populate interstitial macrophages after an initial challenge and may indicate that these cells are responsible for the increased protection we observed. (n=4-10, median plotted, 2-way ANOVA, post-hoc Tukey’s multi-comparison test. P< 0.05 **p<0.01).

The myelomonocytic driver LysM is known to target both macrophage and neutrophil populations, and loss of EZH2 in neutrophils impairs the migratory behavior of these important migratory phagocytes (19). However, the CX3CR1 driver does not affect neutrophils, permitting analysis of the role of EZH2 in macrophages in lung inflammation models in which a neutrophil response is prominent, and essential.

There was a small but significant increase in the total number of cells in targeted mice in broncho-alveolar lavage (BAL) derived from the lungs of LPS exposed mice, which flow cytometry revealed was caused by a significant increase in the number of neutrophils (Cd11b+, Ly6G+) (Fig 4B, Fig S6A). There were no differences in numbers of alveolar macrophages (CD11c +, Siglec F+, Cd64+) or cytokines CXCL1 (Figure 4B), CCL2, IL-6, or IL-1a (Figure S6B).

We next assessed responses to bacterial infection, using an intranasal delivered *Streptococcus pneumoniae* infection model. EZH2-targeted mice were assessed 40h after infection. The numbers of colony forming units (CFU) recovered from lung were not different by genotype, but in marked contrast we observed substantial differences in the blood compartment – with no CFUs detected in blood of targeted mice (Fig 4C). Flow cytometry analysis of the lung digest revealed no difference in the total number of cells/mg of tissue after infection. However, the number of interstitial macrophages in the lung after infection was significantly greater in the EZH2^fl/fl^ Cx3CR1-cre mice (Fig 4D, FigS7). Thus, loss of EZH2 in pulmonary macrophages greatly enhances their ability to restrict the systemic spread of a localised infection. This was a really striking phenotype proving a role for macrophage EZH2 in regulating pulmonary bacterial defence, with a gain of function seen in macrophage-specific EZH2 knockout animals.

### EZH2 deletion in pulmonary epithelial cells does not affect influenza responses

Our data revealed a striking in-vivo phenotype following targeting of macrophages. We have previously shown that pulmonary bronchial epithelial cells play a sentinel role in gating TLR-4 mediated time-of-day responses to LPS and inflammatory responses, regulated by specific elements of the epithelial cell clockwork (20, 21). We therefore targeted EZH2 in bronchial epithelial cells using a CCSP-cre driver (20). Lung tissue of targeted mice was of normal histological appearance, and mice developed normally. However, we did observe a very subtle change in circadian period resulting from either genetic deletion of EZH2, or its pharmacological inhibition (Fig S8A-H). Accordingly, we tested diurnal responses to LPS, and this revealed significant time-of-day differences in the magnitude of neutrophilia and cytokine/chemokine responses, compatible with earlier studies (20), but which were not significantly different in EZH2-targeted mice (Fig S8I-L).

As the bronchial epithelium is targeted by the prevalent influenza virus we tested the responses to influenza inoculation. In both targeted and wild-type animals there was a marked reduction in body weight and establishment of an inflammatory response (Fig5 A-E), but no effect of genotype, nor differences in neutrophil or macrophage cell number. Thus, despite evidence for the importance of the bronchial epithelial cell as a local-acting stem cell population (20) and the role of these cells in gating time-of-day responses to LPS, targeting of EZH2 has no significant impact on the kinetics of inflammatory responses nor response to epithelial-targeted viral infection.

### Neutrophil EZH2 is essential to confer protection against bacterial infection

Previous work has shown that EZH2 is critical for neutrophil extravasation (19). To investigate this phenotype further in our physiological models we went on to test inflammatory responses of LysM-cre.Ezh2^fl/fl^ mice, which lack EZH2 in both macrophage and neutrophil cells. Initially, we tested pulmonary inflammatory challenges to nebulized LPS exposure. BAL analysis at +5h revealed a significant reduction in recruitment of neutrophils, but no significant differences in macrophage number (Fig 6A), as anticipated. We next assessed whether production of pulmonary neutrophil chemo-attractants was disrupted, and analyzed BAL and peripheral blood for LPS-induced expression of the neutrophil chemokine Cxcl5 and the monocyte chemokine Ccl-2. This revealed an increase in CCL2 in BAL from LysM-cre.EZH2 mice following aerosolized LPS challenge but no difference in blood or CXCL5 from either tissue. (Fig 6B,C). Therefore, it appeared that the defect in neutrophil recruitment lay with the neutrophils rather than with a lack of appropriate chemokine signal.

**Figure 5.**
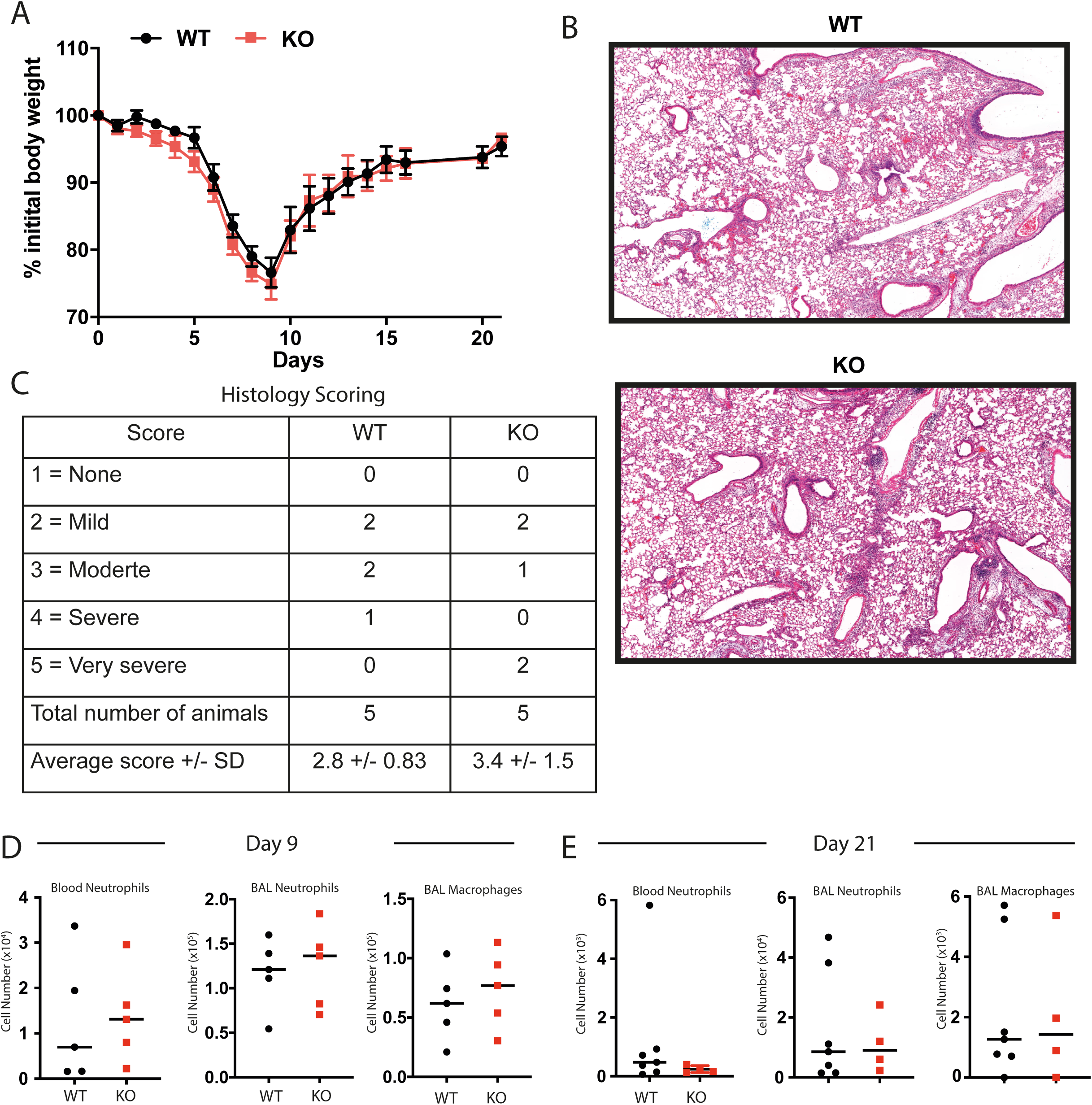
Influenza responses in bronchial epithelial EZH2 null mice. A) EZH2 was disrupted in bronchial epithelium using the CCSP-icre crossed with EZH2^fl/fl^. EZH2^fl/fl^ CCSP-icre (KO) and EZH2^fl/fl^ (WT) mice were infected with influenza, and responses tracked through time by measurement of body weight. The nadir at day 9 was followed by recovery, in both genotypes. Loss of EZH2 in these cells did not alter the trajectory of response to infection. B) Histological analysis of lungs at Day 9 (H and E stain) did not reveal any differences by genotype. C) At the peak of infection peripheral blood, and lung content of immune cell sets did not differ by genotype. D, E) Analysis of blood and lung immune cell sets after recovery did not differ by genotype at day 9 or 21.

**Figure 6:**
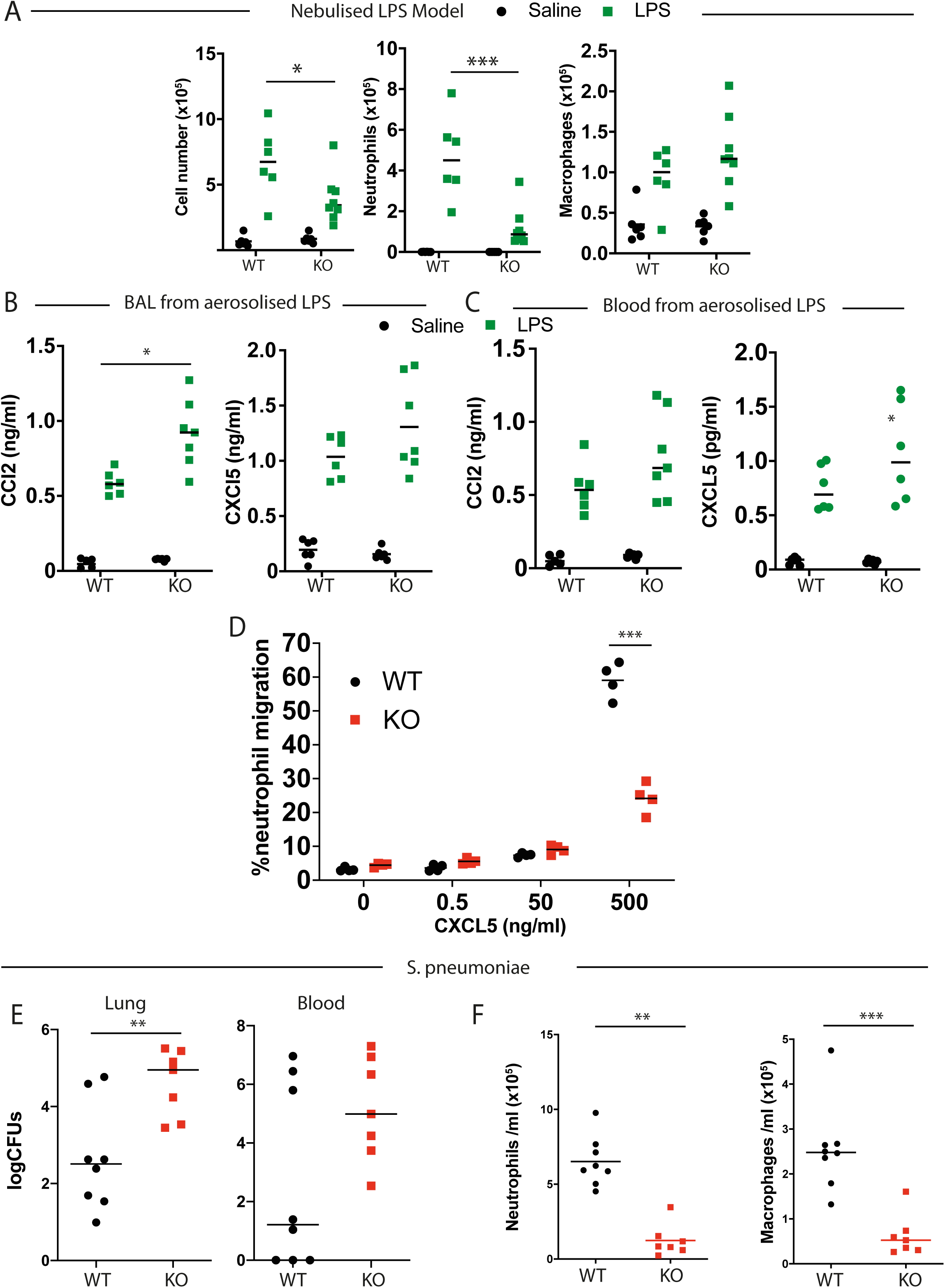
EZH2 differentially impacts myeloid cell types. A) Nebulised LPS was administered to either EZH2^fl/fl^ (WT) or EZH2^fl/fl^.LysM-cre (KO) animals. After 4 hours, lung immune cell content was measured by BAL and FACS analysis. This clearly demonstrated a reduced cellular infiltrate after challenge that is accounted for by a lack of neutrophils, but minimal changes in macrophage numbers. (n=6-8, median plotted, 2-way ANOVA, post-hoc Tukey’s multi-comparison test. *p<0.05, ***p<0.001) B,C Chemokine content from (A) was determined by ELISA. This revealed an intact chemokine response in BAL (B) or blood (C) from EZH2^fl/fl^.LysM-cre mice (n=6-8, median plotted, 2-way ANOVA, post-hoc Tukey’s multi-comparison test. *p<0.05). D) To evaluate behaviour of isolated neutrophils, cells were isolated from the bone marrow of genetically targeted mice and were placed in Boyden chamber across a membrane from a CXCL5 gradient. This clearly demonstrates that neutrophils lacking EZH2 have a significantly reduced capacity to migrate towards that signal. (n=4, median plotted, 2-way ANOVA, post-hoc Tukey’s multi-comparison test. ***p<0.001). E) Mice of the indicated genotype were inoculated with pneumococcus, and 48 hours later the bacterial content of lung tissue, and peripheral blood were measured. Significantly higher CFUs were observed in the lung (n=7-8, median plotted, Mann-Whitney *U*-test **p<0.01). F) Lung immune cell content was measured from BAL by FACS analysis, post-pneumococcal infection. This revealed striking suppression of both neutrophils and macrophages in EZH2 targeted mice. (n=7-8, median plotted, Mann-Whitney *U*-test **p<0.01, ***p<0.001).

To test this we used a modified Boyden chamber to measure neutrophil migration against a CXCL5 gradient. This revealed a marked defect in neutrophil response, supporting a major migratory phenotype likely resulting from the cytoskeletal re-modelling effects of EZH2 loss in these cells (Fig 6D).

To evaluate the biological relevance of this phenotype in the context of bacterial infection, we exposed LysM-cre.EZH2^f/f^ mice to infection with S. pneumoniae. After 48h of infection we observed a significant increase in the bacterial colonisation of the lung, but no change in the recovery of bacteria from lung and peripheral blood (Fig 6E).

Lung tissue was digested and this revealed a significant reduction in total cell counts, both in neutrophil and macrophage populations in the homogenate (Fig 6F). The reduction in neutrophil recruitment was expected, but the attendant loss of macrophages was a surprise, and likely results from the extent of infection, and inflammation.

### Defective neutrophil migration can be rescued by the transplant of EZH2 intact neutrophils

We next set out to define the role of Ezh2 in neutrophil protection from pneumonia by using an adoptive transfer model, in which we restored Ezh2 intact neutrophils to LysM-cre.Ezh2 fl/fl targeted animals. Transferred neutrophils could be tracked by expression CD45.1 (Fig 7A) Following adoptive transfer, animals were exposed to aerosolised LPS.

**Figure 7.**
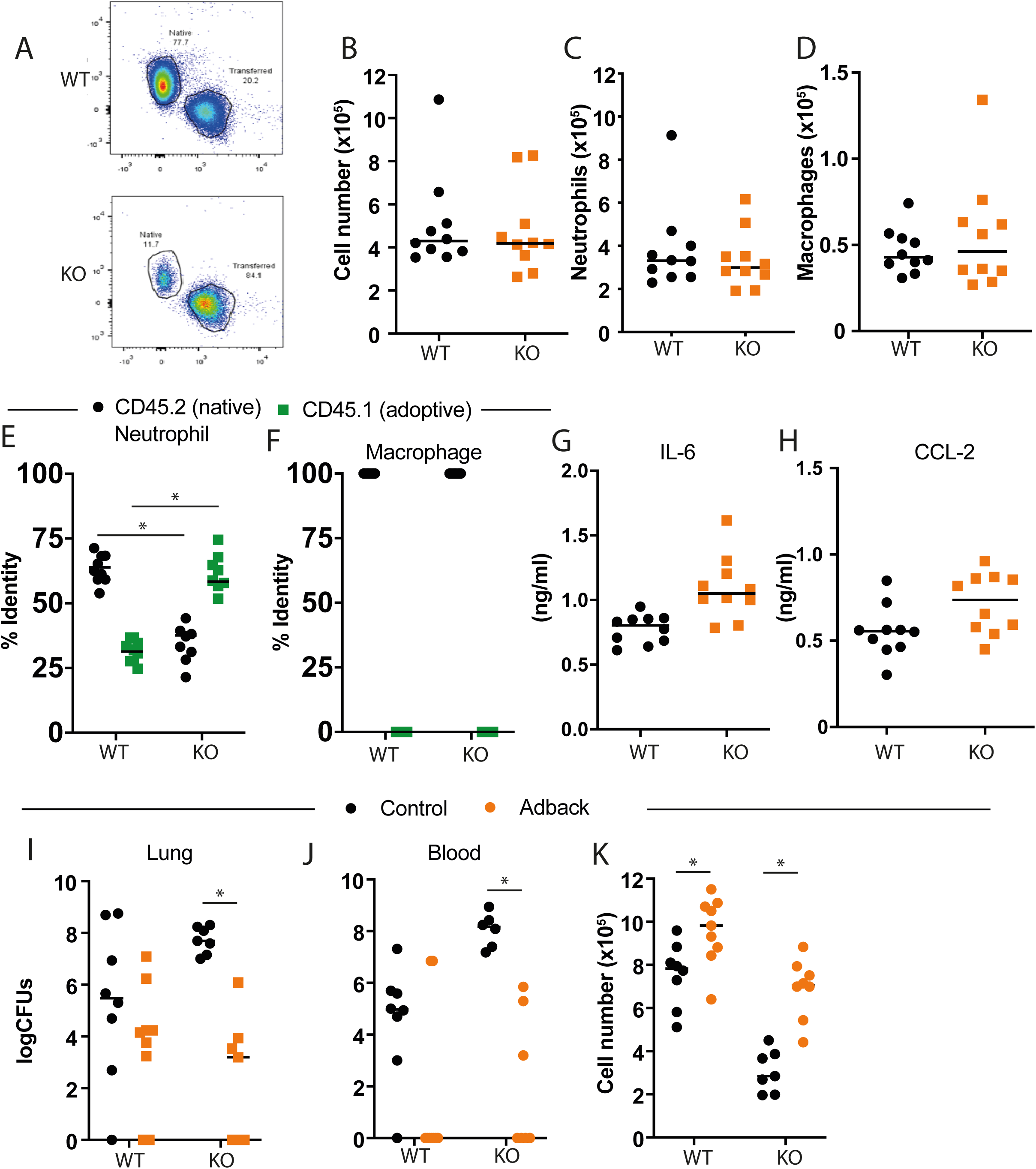
Functional EZH2 in Neutrophils is required for normal immune responses in the lung. A-C) Bone marrow from control animals expressing CD45.1 (PEP3 mice) was used to purify neutrophils which were infused in the EZH2^fl/fl^ (black circle) or EZH2^fl/fl^.LysMcre (red square) animals by tail vein injection. Animals were exposed to nebulized LPS as above, and lung immune cell infiltration in BAL determined by flow cytometry D) representative flow cytometry flowplot. E) The origin of lung neutrophils was measured by flow cytometry analysis of CD45.1 (donor), and CD45.2 (host). F) Lung macrophage origin was also assessed by flow cytometry for CD45.1 status. The adoptive transfer rescued neutrophil recruitment to the lung and there were no adopted macrophages. G,H) Cytokine and chemokine concentration in BAL was measured by ELISA. Stats needed – looks different but is it? (Mann-Whitney U-test; NS p>0.05). I-K) Mice (EZH2^fl/fl^.Lysmcre = KO) treated with adoptive transfer of neutrophils from CD45.1 wild-type animals were compared to wild-type control mice also infused with the same number of neutrophils. Animals were then inoculated with pneumococcus, and the animals were killed and analyzed 48 hours later. The lung bacterial content (I), and peripheral blood bacterial content (J) were determined, and (K) lung neutrophil infiltration assessed by BAL, and flow cytometry analysis. Adoptive transfer of WT neutrophils was sufficient to rescue the normal anti-bacterial phenotype (n=6-9, Kruscal-Wallis with Dunn’s multi-comparison test (I&J), 2-way ANOVA, post-hoc Tukey’s multi-comparison test (K). *p<0.05,).

In marked contrast to our earlier studies, there were no genotype-specific differences in the number of BAL cells after LPS challenge (Fig 7B) or a change in the number of neutrophils (Fig 7C) or alveolar macrophages (Fig 7D).

There was a marked change in the genotype of the neutrophils between the LysM-cre.EZH2^f/f^ and EZH2^f/f^ control animals (Fig 7E). However, all the macrophages were CD45.2, and therefore of host animal origin (Fig 7F). We also noted no genotype-specific differences in concentrations of IL-6 (Fig 7G) or Ccl2 (Fig 7H) in BAL. In order to show that neutrophil Ezh2 was essential to combat bacterial infection of the lung, we used this adoptive transfer model to test responses to intranasal infection with *S. pneumoniae*. Here, we noted a significant reduction in lung bacterial burden of Ezh2^fl/fl^ LysM-cre mice after neutrophil add-back, and no significant difference in the Ezh2^fl/fl^ controls (Fig 7I). We also saw a reduction in bacterial recovery of bacteria in the add-back animals, compared to the LysM-cre.EZH2f/f host (Fig 7J). Further, BAL cell recovery after infection showed a significant increase following neutrophil add-back in both genotypes, with the Ezh2^fl/fl^ LysM-cre mice showing an increase to control levels of BAL neutrophils (Fig 5K). Thus, we demonstrate complex and opposing effects of EZH2 deficiency in monocytic lineage cells, in which we observe an enhanced pro-inflammatory responses of macrophages, while loss of EZH2 in neutrophils impairs migration and responses to bacterial infection.

## Discussion

Regulation of innate immunity at mucosal surfaces is exquisitely sensitive to context, and requires interplay between resident immune surveillance cells, notably macrophages, epithelial cells, and recruited cells, notably neutrophils (20). There has been interest in targeting epigenetic mechanisms to regulate the function of some haematopoeitic cell malignancies, especially using drugs against the polycomb II complex (15). In addition, some reports suggest targeting the enzymatic core of polycomb II, EZH2, impacts on immune responses in the non-malignant state (14). Therefore, it is timely to investigate the role of EZH2 in mucosal immunity.

Here, we investigate the role of EZH2 in pulmonary mucosal inflammation. We used inflammatory challenge models targeting the lung, including nebulized lipopolysaccharide, (LPS), and also two pulmonary infectious diseases (influenza A and Pneumococcus). Initially, we targeted EZH2 in myeloid-lineage cells using a LysM-cre driver crossed to a floxed EZH2 line of mice, and showed no significant change in cell-surface markers for EZH2 deletion. Loss of EZH2 or pharmacological inhibition using a highly-specific, SAM-binding site compound (15) elicited pro-inflammatory cytokine responses in cultured macrophage cells and in-vivo in the lung. We next showed that EZH2 has specific and selective action inhibitingTLR4-mediated pro-inflammatory pathways. Since LysM targets both neutrophil and macrophage populations, we switched to use of a macrophage-specific CX3CR1-cre line, and tested responses to Pneumococcal infection. Despite similar bacterial loads in the lung, loss of EZH2 completely protected animals from septicaemia, leading us to conclude that loss of EZH2 in pulmonary macrophages greatly enhances their ability to restrict the systemic spread of a localized infection.

We extended these studies to an influenza infection model, which is known to target pulmonary epithelial cells and to depend on lymphocytes and anti-viral signalling molecules including the type I and II interferons (22). Loss of EZH2 in bronchial-epithelial cells did not alter the trajectory of the disease response, supporting our earlier observations that the primary role of EZH2 is via a TLR4 mediated pathway.

We returned to the pneumococcal infection model, and extended loss of EZH2 to neutrophils by using Ezh2 fl/fl LysM-cre mice. Mice were exposed to LPS, either by injection to the peritoneal cavity, or nebulised to the lung. Despite modest increases in inflammatory cytokines and chemokines there was a loss of neutrophil recruitment with myeloid loss of EZH2. This resulted from impaired neutrophil migratory capability, as demonstrated using Boyden chamber studies. Pneumococcal infection of the EZH2fl/fl LysM-cre animals resulted in increased bacterial proliferation in the lung, associated with impaired recruitment, or retention of macrophages and neutrophils. However, transfusion of EZH2 intact neutrophils was sufficient to rescue the pneumococcal response, isolating the defective response to the neutrophil cells.

Examining macrophages isolated from different compartments we identified a gain in inflammatory response to canonical TLR activation. We were able to show that the effect of the EZH2 inhibitor UNC1999 was entirely dependent on the presence of EZH2, and thus confirmed the exquisite specificity of the tool compound. Our findings were compared to those in previous reports, and although we could confirm the specific time and LPS-concentration specific reduction in KC response with loss of EZH2 the majority of the cytokines and chemokines we profiled showed an increase in response to activation resulting from EZH2 loss of function (14). To investigate the basis for this change we investigated gene expression in macrophages lacking EZH2. Here we discovered increased expression of TLR4 signaling component genes, and using a transcription factor activation assay identified a specific impact of EZH2 on the TLR4 cascade, with no impact on the viral TLR3 response, which led us to the hypothesis that the EZH2 effect is likely specific to particular immune stimuli. This specificity is also likely the explanation of the differences reported here compared to those in the earlier work (14). Extension of the mechanistic studies to RNA-SEQ profiling revealed a broad similarity in terms of upstream regulatory pathways dependent on EZH2, but there was significant enrichment in increased predicted activity of PPARg, and in contrast a predicted decrease in the activity of Tcf7, Id3 and meis1. Integration of gene expression with predicted activity based on target gene changes revealed that the IRFs1, and 7, and NFkB were seen to show an increase in both expression, and function. These changes all point to a gain in innate immune response.

In order to distinguish macrophage effects from those of the neutrophils we moved to develop new lines of mice with CX3CR1-cre targeting EZH2 deletion. This approach allows in-vivo testing of innate responses, with neutrophil responses as part of the output. Again, we compared macrophages from different compartments, and here we discovered that peritoneal macrophages gained a significant new population of cells with low MerTK. We speculated that these arose in the peritoneum as a result of altered macrophage development, but efforts to identify emergence of a low MerTK population in differentiating bone marrow cells failed. Therefore, the origin of the MerTK low cells remains obscure, but was confined to the peritoneal cavity. In contrast we could not detect any phenotypic differences in the alveolar macrophage population, and so we moved to test pulmonary immune responses in-vivo. We started with nebulized LPS, to extend the earlier in-vitro analyses. Here we saw modest increase in neutrophil infiltration resulting from EZH2 loss, but overall the impact was surprisingly modest. However, LPS is a mimic of gram-negative bacterial infection, which is less prevalent in the lung. So, we switched to infection with the Gram-positive bacterium pneumococcus, a prevalent pulmonary pathogen, and the cause of massive human illness burden. These studies identified a very major gain in immune response, with animals lacking EZH2 just in the macrophages, having complete protection from septicemia (extension of the pneumococcal infection to the bloodstream). Anti-bacterial gain was associated with increased numbers of both alveolar and interstitial macrophages. The gain in function here is far greater than anticipated from the LPS responses, and the gain in cell numbers suggests a more profound change in macrophage behavior resulting from EZH2 loss, possibly extending to reduced cell death in response to infection. As can be seen infection of control animals results in loss of macrophage numbers, a well-described response, but this is reversed in the absence of EZH2.

The focus on lung macrophages should not overlook the role of respiratory epithelial cells in regulating responses to pathogens. We deleted EZH2 in bronchial epithelial cells using the well-characterized CCSP-cre driver. Testing LPS responses we did not discover any differences in neutrophilic inflammatory responses in the epithelial EZH2-null animals, but we did, again, see that the time of day effect was marked. This time of day gating is dependent on the operation of a circadian clock in the epithelial cells, a machinery that may be impacted by EZH2. Indeed, we were able to find a small impact of EZH2 on circadian period length when isolated macrophages were studied in-vitro. Although we have previous studied ciliated and non-ciliated bronchial epithelial cells in-vitro these approaches change the cellular phenotype making sustained analysis of circadian parameters unreliable (23). We went on to test influenza infection, again a highly prevalent human lung pathogen, and one which targets the epithelial cell. Here we saw no alteration in the responses to infection, with a similar time course of weight loss and recovery in both genotypes, with very similar lung histology appearances at the height of disease.

As EZH2 played such a major role in the macrophage response to bacterial infection we extended these studies to include the neutrophil, in which EZH2 is reported to exert a non-genomic role in cytoskeletal function, and motility. To achieve this goal we switched back to the LysM-cre driver, which hits both macrophages and neutrophils. We find that the LysM-cre targeting strategy resulted in reduced neutrophil recruitment. This is consistent with previous reports. However, as we have seen in earlier studies simple TLR agonist challenges are not predictive of responses to infections with prevalent respiratory pathogens. Therefore, we moved to pneumococcal challenge again. Here we saw the opposite phenomenon to that in the macrophage-selective EZH2 loss, with impaired protection against infection, and despite increased concentrations of inflammatory cytokines, and monocyte chemokines (CCL2), reduced numbers of macrophages and neutrophils in the lung. Isolated neutrophils did indeed fail to migrate towards a chemokine source, confirming the migratory deficit in these manipulated cells (19). In contrast the earlier studies here, the loss of macrophage numbers likely results from apoptosis resulting from excess bacteria, a well-characterized response to bacterial sepsis in the lung (24)

In order to investigate the contribution of the neutrophil deficit to the pneumococcal response in LysM-cre targeted mice we used a neutrophil adoptive transfer approach. Here, we used a tail vein injection of bone marrow sourced neutrophils from wild-type mice to both control, and LysM-cre targeted EZH2 null animals. The origin of the immune cells was tracked by using variant CD45 expression, and FACS. Here we can see no transfer of macrophages, but marked gain of transfused neutrophils in both recipient genotypes. The transfused neutrophils were sufficient to restore the anti-bacterial response in the LysM-cre mice, so confirming that the defect in neutrophil recruitment to the lung was responsible, and not a residual impact on macrophage function. A likely explanation for this is due to protein methyltrasferase-mediated activity of EZH2 on talin-1 cleavage, which disrupts binding to F-actin, leading to disrupted adhesion complex formation and aberrant cell migration (19). This is also consistent with more recent studies showing that disruption of talin1 methylation sites impairs transmigration of neutrophils across the peritoneal vascular epithelium during inflammation, an effect also mimicked by treatment with an EZH2 inhibitor (25).

In summary, we have characterized the role of EZH2 in myeloid cells during inflammatory challenges. EZH2 in macrophages plays a specific role in controlling a TLR-4:NF-kB circuit. This results in increased macrophage cytokine production, greater neutrophilia in response to inflammatory challenges and protection in in vivo pneumococcal challenges. As EZH2 inhibitors are currently under evaluation for the treatment of various cancers, our study suggests caution, due to the immune consequences of targeting EZH2. Furthermore, the exquisite specificity of immune response to EZH2 manipulation raised the possibility that pulmonary neutrophilic diseases, and processes, including ARDS, may benefit from pulmonary delivery of EZH2 inhibitors.

## Acknowledgements

The work is supported by Biotechnology and Biological Sciences Research Council BBSRC grants awarded to ASL and DWR (BB/L000954/1, BB/K003097/1), and an MRC programme grant MR/P023576/1. DWR and ASL are Wellcome Investigators, Wellcome Trust (107849/Z/15/Z). GBK was supported by an MRC Clinical Research Training Fellowship (MR/N002024/1) and is currently an NIHR Academic Clinical Lecturer. TH was supported by an MRC PhD training award co-supervised by Kathryn Else. We are grateful to Alexander Tarakhovsky, The Rockefeller University for the Gift of the EZH2 murine driver and also to Jian Jin, from Mount Sinai for the gift of UNC1999. We thank Andy Hayes, Michal Smiga, Leo Zeef and I-Hsuan Lin, (Bioinformatics and Genomic Technologies Core Facilities, University of Manchester), for providing support with regards to the NanoString and RNA-seq analysis. Ryan Vonslow for technical support. Pete Walker (Histology facility, University of Manchester) for histology support. We would also like to acknowledge the BSF staff, University of Manchester, for animal care.

## METHODS

### Animal Studies

All experimental procedures were carried out in accordance with the Animals (Scientific Procedures) Act of 1986. Male C57BL/6J mice aged 8-12 weeks were purchased from Harlan Laboratories, UK. EZH2flox/flox mice possess loxP sites upstream and downstream of the 2.7kb SET domain and were a gift from Alexander Tarakhovsky (Rockefeller, New York). These mice were bred with LysMcre/+, a cre recombinase under the control of lysozyme M, to target Ezh2 for deletion in myeloid cells, including macrophages and neutrophils. Offspring produced were either EZH2flox/flox with no LysM-cre present, these mice acted as littermate controls, or were positive for the cre driver. In order to genotype these mice and elucidate the correct loxP sites had been inserted the following primers were used for EZH2:

Reverse 1: 3’of loxp: 5’-AGG GCA TCA GCC TGG CTG TA-3’

Forward 2: 5’ of loxp: 5’-TTA TTC ATA GAG CCA CCT GG-3’

Forward 3: left loxp 5: -ACG AAA CAG CTC CAG ATT CAG GG-3’

Primers were used at a final concentration of 2.5μM and were genotyped using the GoTaq Hot Start Polymerase (Promega). The PCR product produced using these primers was run on a 2% agarose gel at 45 volts for 3h and mice homozygous for the loxP sites presented a band at 731bp and those wild type for loxP expression with a band at 691bp. Mice homozygous for the flox were selected and genotyped for the expression of LysM-cre using primers diluted to 2.5μM:

Forward: 5’ – CTTGGGCTGCCAGAATTCTC – 3’

Reverse: 5’ – CCCAGAAATGCCAGATTACG – 3’

The presence of the LysM-cre gene was confirmed by a band at 700bp on a 2% agarose gel for 45 minutes at 80 volts.

To target macrophages exclusively and avoid broad myeloid targeting, EZH2flox/flox mice were bred with CX3CR1cre/+ mice. EZH2 genotyping was performed as stated. CX3CR1 cre/+ presence was determined using primers diluted to 2.5μM:

Forward: 5’ - CCC CAA TGT TCA ACA TTT GC – 3’

Reverse: 5’- GGA CTG GGG ATG TGG GAG – 3’

cre postivie mice were determined by the presence of a band at 500bp following the PCR product being run on a 2% agarose gel at 80 volts for 45 minutes.

For investigations into the role of EZH2 in airway epithelium cells mice possessing the cre driver CCSPicre were crossed with mice possessing the EZH2flox/flox allele. The presence of loxP sites in EZH2 was confirmed as before. CCSPicre was evaluated using primers at 2.5μM:

Forward: 5’ – TCTGATGAAGTCAGGAAGAACC – 3’

Reverse: 5’ – GAGATGTCCTTCACTCTGATT – 3’

and mice were selected as cre positive if a band was identified at 500bp following the PCR product being run on a 2% agarose gel at 80 volts for 45 minutes.

Evaluation of the role of Bmal in cells of the myeloid lineage Bmalfl/fl mice were crossed with mice possessing the LysM-cre driver. The presence of LysM-cre was determined as described. Bmal was genotyped

Finally, the use of the Per2::Luc transgene was measured using 2.5μM primers:

Common: 5’ – CTGTGTTTACTGCGAGAGT – 3’

WT Reverse: 5’ – GGGTCCATGTGATTAGAAAC – 3’

Mutant Reverse: 5’ – TAAAACCGGGAGGTAGATGAGA – 3’

The PCR product was run on a 2% agarose gel for 40 minutes at 80volts. A WT band at 230bp or a mutant band at 650bp showed the absence of presence of the Per2::Luc construct.

### Bone marrow derived macrophages (BMDM)

C57BL/6 mice are obtained and sacrificed via cervical dislocation. The hind legs were removed, the muscle and skin dislodged using gauze and to clean the bone before being placed in ice cold sterile dulbecco’s phosphate buffered saline (PBS), this procedure was repeated with the fore legs. The bones were then placed in ice-cold DMEM (Sigma) with 10% foetal bovine serum (FBS) (Life technologies) and 1% penicillin/streptomycin (Sigma) in a primary tissue culture hood. The ends of the bones were removed and a 17-gauge needle was used to flush out the bone marrow using the culture media. The bone marrow was disrupted and centrifuged at 1600rpm for 6 minutes and the supernatant removed. Cells were re-suspended in 5ml culture medium plus macrophage colony stimulation factor (M-CSF) (Affymetrix eBioscience) at a concentration of 50ng/ml and counted on a Nucleocounter NC-250. Cells were then plated into either 6-well plates at a concentration of 1.5×106, 12 well plates at 0.8×106 or 24-well plates at 0.4×106/ml and covered with 1.5ml of culture media + M-CSF. Cells were kept in a primary incubator at 37°C and 5% CO2 unless otherwise stated and media was changed every 3 days by washing in PBS and replacing culture medium.

### PECS

Cells were isolated by washing the peritoneal cavity with ice cold PBS several times and collecting the washout. This was then centrifuged at 200G for 5 minutes at 4°C and the cells re-suspended in DMEM/10% FBS/1%penicillin/streptomycin. Cells were then counted and plated at a concentration of 1×106/ml in 6-well plates for 24h. Non-adherent cells were washed away with pre-warmed (37°C) DMEM. The remaining adherent cells were termed peritoneal macrophages.

### Alveolar Macrophages

Cells were purified using by culling a mouse by cervical dislocation, with care taken to avoid compromising the trachea. A 23 gauge needle was used to pierce the trachea and fill the lungs with 1ml ice-cold BAL fluid (PBS, 1% FBS/0.5mM EDTA). BAL fluid was pulled back from the lungs and placed in an Eppendorf on ice. Eppendorfs were centrifuged at 200G for 5 minutes at 4°C and the pellet was resuspended in DMEM (10% FBS/1% penicillin-streptomycin). Cells were plated out in 24-well plates and cells were left to adhere for 24h before the removal of non-adherent cells by washing in pre-warmed (37°C) DMEM. The remaining adherent cells were termed alveolar macrophages.

### Neutrophil isolation

Hind limbs were isolated as above. The bone marrow that was flushed through and was counted and resuspended at 1×107/ml with 5% normal rat serum (EasySep) in PBS and was incubated on ice for 15 minutes. Mouse Neutrophil Enrichment Cocktail (EasySep) containing primary antibodies against non-neutrophils was added at 100μl/ml and incubated on ice for 15 minutes. Bone marrow was washed in 10ml of pbs, centrifuged at 8000rpm for 5 minutes and resuspended at 1×107/ml in pbs. Magnetic secondary antibodies were added at 100ul/ml, vortexed and placed on ice for 15 minutes. The sample was placed in a magnetic field for 2 minutes at room temperature and the remaining PBS containing the neutrophils was removed.

### Adoptive transfer

Neutrophils were isolated, as described above, from PEP3 mice that possess a naturally occurring CD45.1 isoform as opposed to CD45.2 found in C57BL/6 mice. These neutrophils were then diluted to contain 5×106 neutrophils/ml in saline. Target mice were kept in a heat box at 37°C for 20 minutes and were then restrained. These mice then had 200μl of neutrophil solution containing 1×106 neutrophils injected into the tail vein before being returned to the home cage. Mice were left for 1h to allow neutrophil circulation prior to any experiments taking place.

### Intra-peritoneal LPS

LPS (Sigma) was reconstituted in saline and mice were injected using a 23 gauge needle at a dose of 1mg/KG in 200ul. After injections, mice were returned to their home cages to generate an immune response in the peritoneal cavity for a time period stated in each individual experiment. Macrophage activity was evaluated by isolating the cells from the peritoneal cavity as described.

### Aerosolised LPS

Mice were placed in individual restraining compartments of an inExpose chamber (Scireq). The lid of the compartment was then securely placed on the chamber, placed in a fume hood and a nebulisation unit (Scireq) was attached to the chamber. Saline or LPS reconstituted in saline (2mg/ml) and was then placed in the nebulisation unit. The flow rate of the machine was set to 4L/minute and the nebuliser was turned on to aerosolise the LPS. Mice were then exposed to the aerosolised LPS or saline control for twenty minutes. Mice were then returned to their cages for 5 or 24h before being culled by intra-peritoneal overdose of pento-barbitone or cervical dislocation.

### Broncho-alveolar lavage

After mice were euthanised, the trachea was exposed by blunt dissection and a 23-gauge needle, connected to a 1ml syringe filled with BAL fluid, was inserted into the trachea. The BAL fluid was then forced into the lungs by pressure and was left for 1 minute. BAL fluid was then extracted from the lungs and then fluid was placed in an Eppendorf on ice.

### Cell counting

Cells were pelleted after in vivo experiments and were resuspended in 1ml PBS. Acridine Orange provides a total cell count whilst DAPI provided information on the total number of dead cells allowing the calculation of viability of each sample using NucleoView software for Nucleocounter.

### Exposure to Streptococcus Pneumoniae

Streptococcus pneumoniae strain D39 (National collection of Type Cultures 7466) was prepared in Todd Hewitt Broth (Sigma) and incubated at 37°C 5% CO2 until an optical density of OD600 was achieved from 10ml of growth media. Mice were then placed under light anaesthesia using isoflurane. Whilst under light anaesthesia the mice were scruffed and, using a pipette, 100μl of *S. pneumoniae* was dropped onto the nose to intranasally administer 2×10^4^ colony forming units of D39. Mice were closely monitored and weighed every 12h to assess health (a threshold of 20% weight loss was used as an end-point) and were culled 48h after exposure by intra peritoneal pentobarbitone overdose in a category 2 fume hood. To evaluate the infective burden 100μl of peripheral blood was obtained immediately after cull and placed in 5μl heparin (Sigma) to prevent clotting and was placed on ice. The chest cavity was then opened and the lungs were exposed. All lobes of the lung were removed and placed in ice-cold DMEM and placed on ice.

### Culture of *S. pneumoniae*

To understand the infective burden in each animal blood and lung tissue were obtained as above. Lung tissue was digested as described in section 2.15. Serial dilutions of blood and lung tissue were then made in 96-well plates. 20μl of the serial dilution of blood or lung digest was then placed onto horse-blood agar plates in duplicate for each sample. Blood agar plates were produced by making a 3.9% Columbia blood agar base (Sigma) in deionised water and autoclaving. Agar was allowed to equilibrate to 50°C in a water bath before 5% defibrillated horse blood (OXOID) (TCS Biosciences) was added. The solution was then poured into 10cm plates and allowed to set at 4°C. Once bacteria was added to the agar plates they were left to air-dry in a cell culture hood for 30 minutes before being placed in a 37°C incubator at 5%CO2 for 24h. Blood agar plates were then removed from the incubators and the bacterial growth was counted at each serial dilution to understand the bacterial growth in the blood and lung of each mouse.

### Evaluation of bioluminescent rhythms

To analyse the rhythms of the Per2 protein peritoneal macrophages were obtained, as described, from mice with the Per2::Luc construct. Macrophages were grown in a 25-mm culture dish and were synchronised using 48h of temperature oscillations. After synchronisation the culture media was replaced with 1ml recording media (recipe in appendix). Vacuum grease (Dow Corning) was placed on the rime of the cell culture dish and a 50mm glass coverslip was placed on the top of the grease and gentle pressure applied. The dish and coverslip were placed in a 37°C incubator containing a photon multiplier tube (PMT – H6240 MODI, Hamamatsu Photonics, Shizuoka, Japan). Multiple PMTs existed within an incubator in either an 8 or 16 channel set up. Recordings were taken every 60s of bioluminescence.

### Rhythm analysis

Rhythms of bioluminescence were recorded as photon counts/minute and normalised to a 24h moving average to produce oscillations about a mesor of 0counts per minute. Normalised data was analysed using cosinor analysis from Biodare 2 (Zielinski et al., 2014).

### Lung digestion

The left lobe of the lung was placed into a bijou containing ice-cold DMEM during the cull. DMEM was removed and the lung lobe was then chopped into small sections using dissection scissors. 1ml of digestion media: DMEM containing 2.7% Liberase (Roche) and 0.05% DNase I (Roche) was added to each bijou and they were placed in a shaking incubator at 37°C for 30 minutes. To stop the digestion 0.005mM EDTA in DMEM was added to the digestion media in an equal volume. The digestion was then forced through a 70um culture sieve. At this point samples required for S. pneumoniae infection analysis were taken and placed on blood agar plates. The homogenate was then centrifuged at 1600rpm rpm for 6 minutes at 4°C. The pellet we re-suspended in 2ml red-blood cell lysis buffer (Roche) for 4 minutes before the addition of 10ml DMEM to stop cell lysis. The solution was centrifuged again and red-blood cell lysis was repeated if the pellet was not mostly white in colour. This pellet was then available for cell counting and flow cytometry.

### RNA extraction from tissue

RNA was isolated from tissues using Trizol reagent (Sigma).

### RNA analysis

RNA was isolated from cells using the RNeasy mini kit (Qiagen).

### Reverse Transcription

All samples of RNA were treated with 1μl of RQ1 DNase (Promega). DNase treated RNA was converted to cDNA using the RNA-cDNA kit (Applied Biosystems) in accordance with the manufacturer’s instructions.

### Quantitative Real-time PCR (qPCR)

cDNA was analyzed in 96-well MicroAmp optical reaction plates (Life Technologies). All reactions were performed in a StepOnePlus system (Life Technologies). Relative expression of genes was determined using the threshold cycle comparative CT. Gapdh was used as a housekeeping gene and gene expression was normalised to the housekeeping gene and relative quantification values were calculated in relation to a reference sample, which is denote in each experiment. Fold change was determined using the formula: 2-Δ ΔCT = 2-(ΔCt (experimental sample) – ΔCt(control)). Samples were run in either triplicate or duplicate

### Histone purification

Histones were purified from cells by growing macrophages to a density of 3×10^7^ (Active Motif - 40025). Histone H3K27me3 conversion was measured using the Active Motif kit (Epiquik Histone Methyltransferase activity assay kit (P-3005).

### MTT Viability assay

Cell viability after treatment with EZH2 inhibitors was determined using an MTT assay.

### Enzyme-linked immunosorbent assay (ELISA)

The concentration of multiple proteins was determined using ELISA in a variety of samples including BAL, peripheral serum and lung homogenate. All ELISA kits were DuoSet and were obtained from R&D systems, and run according to the manufacturer’s instructions.

### Nanostring

RNA was then run on a Nanostring nCounterTM (Nanostring Technologies) in conjunction with the pre-made CodeSet called Mouse inflammation panel V2 (Nanostring Technologies).

### RNA-seq analysis

The integrity and concentration of the RNA was evaluated on a nanodrop and RNA concentration was adjusted to 20ng/μl in 5μl of endonuclease-free water. The samples were then run on an Illumina Hiseq 4000 (Illumina). RNASeq data was analysed by the University of Manchester Core Facility using strand-specific RNA-seq libraries prepared using the Illumina workflow. The fastq files generated by Illumina HiSeq 4000 platform were analysed with FastQC and any low quality reads and contaminated barcodes were trimmed with Trimmomactic (Bolger, Lohse and Usadel, 2014). All libraries were aligned to GRCm38.p2 (mm10) assembly of mouse genome using Tophat-2.1.0 (D. Kim et al., 2013) and only matches with the best score were reported for each read. The mapped reads were counted by genes with HTSeq (Anders, Pyl and Huber, 2015) against gencode.vM2.annotation.gtf. Differentially expressed (DE) genes were identified by comparing between the treatment groups with DESeq2 (Love, Huber and Anders, 2014). The functional analysis was carried out with R packages topGO (Alexa, Rahnenfuhrer and Lengauer, 2006) and RontoTools (Ansari et al., 2016). This produced a set of differentially expressed genes between each experimental condition. Expression of genes was considered significantly different using a log fold change cut of 2 or -2 and an adjusted P value <0.05. These differentially expressed genes were then analysed using a KEGG ontology pathway analysis to determine the molecular pathways that contained differentially expressed genes altered by the loss of Ezh2. Motif analysis was performed using the web tool i-cisTarget to identify transcription factor motifs with 10Kb upstream and downstream of the TSS differentially expressed genes after LPS treatment of Ezh2 deficient macrophages compared to LPS treated controls (Imrichová et al., 2015)(http://gbiomed.kuleuven.be/apps/lcb/i-cisTarget.). Upstream regulator analysis was performed using IPA (Qiagen)

### Transcription factor activation assays (TransAM)

RelA and IRF-3 transcription factor activation assays (Active Motif) were carried out using the same protocol with the only difference being the 96-well plates and the antibodies used, each of which corresponding to the respective kit, Rela: 40096, IRF-3: 48396.

### Immunocytochemistry

Bone marrow derived macrophages were isolated as described and were grown on glass coverslips in 12-well culture plates for 7 days. Individual coverslips were then incubated with blocking buffer (2% FBS and 0.01%Tritonx100) for 30 minutes at room temperature before being washed in PBS. Primary antibodies (Histone H3 – Abcam ab1791, H3k27me3 – Active Motif – 39155 and TLR-4 -Abcam 76B357.1) were then diluted 1:1000 in blocking buffer and were incubated on a rocker overnight at 4C. Coverslips were washed 3 times in PBS and secondary antibodies (Oregon Green - Thermofisher O-6382, Texas Red – Thermofisher T-6390) diluted in blocking buffer at 1:500 and applied for 2h in the dark at room temperature. Coverslips were again thoroughly washed in PBS and were mounted using vectamount (Vector Labs) and slides. Images were acquired using Zen software in conjunction with a Zeiss axio imager A1. The quantification of H3 and H3K27me3 staining was performed using ImageJ software (v1.41) following online instructions obtained at https://imagej.nih.gov/ij/docs/examples/ (last accessed 21/03/2018) > Analysing fluorescence microscopy images with ImageJ. The intensity value for H3K27me3 was divided by the intensity value for H3 (control) to determine the intensity of H3K27me3 relative to that of H3.

### Pharmaceuticals

GSK343 (Selleckchem) was resuspended in DMSO and cells were treated at a concentration of 200nm, GSK126 (Sigma) was resuspended in DMSO and cells were treated at 200nm. UNC1999 was used in both in vitro and in vivo studies. For in vitro dosing UNC1999 was re-constituted in DMSO and was used on cells at 2uM concentrations. For in vivo work UNC1999 was reconstituted in vehicle (0.5% of sodium carboxymethylcellulose and 0.1% of Tween 80 in sterile water) using constant agitation for 60 minutes to dissolve the powdered drug. Re-constituted UNC1999 or vehicle was administered to mice by oral gavage at a dose of 50mg/kg in 100μl twice daily for 48h prior to immunological challenge.

### Flow cytometry

1×10^6^ cells were placed in a 96-well plate and cells were suspended in 50μl PBS containing Zombie UV live:dead stain at 1:2000 (Biolegned) for 15 minutes in the dark at RT. Live:dead staining was halted by adding 150μl FACS buffer (PBS/4% FBS) and centrifuging at 2000rpm for 2 minutes at 4°C. Supernatant was flicked off and cells were washed in 200μl FACs buffer and centrifuged as stated. The pellet was re-suspended in 50μl fc-block (CD16/CD32 antibody at 1:50 in FACS buffer) and incubated at RT for 10 minutes. 150μl of FACS buffer was then added to each well and the plate was centrifuged as before. The pellet was then resuspended 50μl primary antibody cocktail (Table 2) or FMO cocktails for 25 minutes in the dark. 150μl FACS buffer was added to each well and plates were centrifuged and then washed three times. If a secondary antibody was required, it was added to FACS buffer 1:5000 and the pellet resuspended in 50μl of FACs buffer with secondary antibody for 25 minutes at RT in the dark. Plates were then washed three times before the addition of 50μl 1% formaldehyde in FACS buffer for 5 minutes at RT in the dark. Plates were then washed twice in FACs buffer and the pellet re-suspended in 200μl FACs buffer. The samples were then acquired on either a BD Fortessa or BD LSR-II using BD-FACSDivaTM software (BD Biosciences).

Flourophore Antibody target Dilution factor FITC Ly6G 200 A647 MerTK-Bio 100 AF700 Ly6C 200 APC/eFluor780 CD3 100 APC/eFluor780 CD19 200 APC/eFluor780 NK1.1 200 APC/eFluor780 Ter119 200 BV421 CD45.2 200 BV605 CD11c 800 BV650 CD45.1/CD45 200 BV711 CD11b 600 BV785 F4/80 200 Live/Dead Blue L/D Blue 2000 PE/CF594 Siglec F 400 PE/Cy5 MHC-II 10,000 PE/Cy7 CD64 100

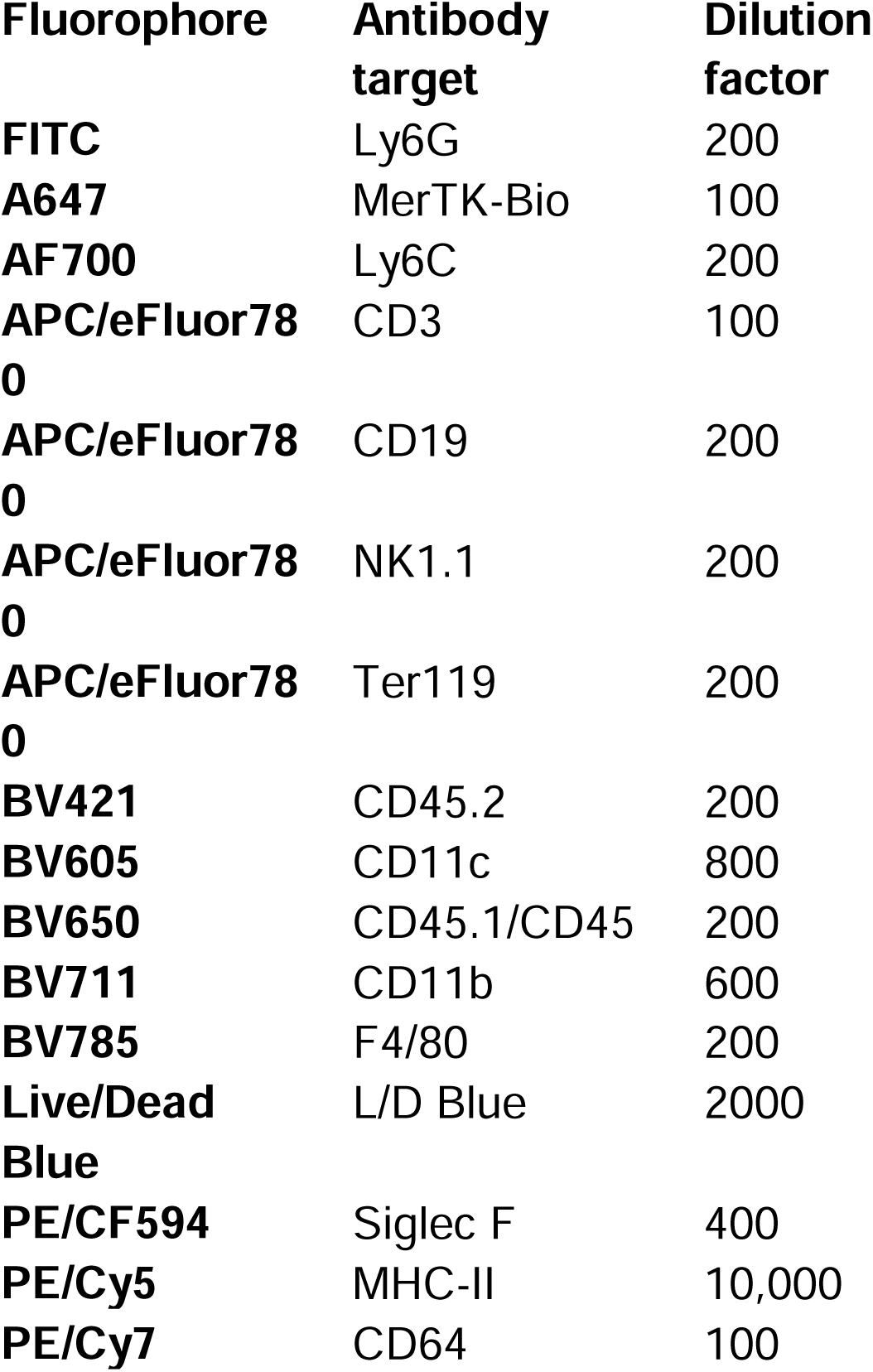

In order to set up the flow cytometers correctly compensation beads appropriate for the panel being analyzed were generated. Compensation beads (BD biosciences) were incubated with each individual primary antibody at the same concentration used on the cells for 10 minutes at RT in the dark. Beads were pelleted by centrifugation at 100G for 5 minutes. The pellet was re-suspended in 200μl of FACs buffer before acquisition. For Live:dead compensation ArC reactive beads (Thermo Fischer) were used according to the manufacturer’s instructions with 1:2000 Zombie:UV Live:dead dye. Immune cells were identified using FLOWJOTM software and all gates were drawn based on published papers using fluorescence minus one (FMO) controls.

### Phagocytosis assay

Cells were cultured as stated and then synchronised. The cells were then detached by washing in room temperature PBS three times and then incubation at 4°C with ice cold 5mM EDTA (Sigma) in PBS for 10 minutes. Cells were then detached with a rubber policeman and checked for health. Cells were then incubated at 37°C with 1mg/ml FITC-DEXTRAN (Sigma) for 1h, control cells were incubated at 4°C to prevent phagocytosis.

The incubation was stopped with ice cold PBS. All cells were then stained for flow cytometry as above. Phagocytosis is expressed as the proportion of F4/80+, cd11b+ cells that have taken up FITC-DEXTRAN.

### Macrophage challenge in vitro

Cells were treated with inflammatory compounds IFN-y (100ng/ml) (Life Technologies), LPS (10ng/ml) or IL-1b (5ng/ml) (R&D systems) or with ‘anti-inflammatory’ compound IL-4 (10ng/ml) (Life Technologies). Compounds were resuspended in sterile water and were added to DMEM for different lengths of time stated in each experiment.

### Boyden Chamber Assay

A Neuro Probe AP48 48-well Boyden chamber (Neuro Probe) was used for all assays. The chamber was prepared by adding 26μl pre-warmed (37°C) CXCL-5 (0, 0.5, 50, 500ng/ml) or CCL2 (0, 0.5, 50, 500ng/ml) (R&D systems) in DMEM to the lower chamber. A filter membrane of either 3μM (neutrophils) or 5μM (Macrophages) pore size (Neuroprobe) was then placed on top of the lower chamber. The silicone gasket and the upper plate were then placed on top of the membrane and the membranes were sealed together with a thumb nuts. 50,000 cells of interest in 50μl of DMEM were added to the top chamber avoiding air bubbles. The Boyden chamber was then placed in an incubator at 37°C and 5% CO2 for 30 minute or 2h. The media was then removed from the top and bottom chambers and the number of cells was counted to obtain a percentage of cells which migrated into the bottom chamber.

### Cell synchronisation

Macrophages were synchronized using a temperature cycling incubator. Culture dishes were placed in the incubator and the temperature changed every 12h from 36.5°C to 38.5°C and then back again. Cells were kept in these conditions for 48h. All synchronizing experiments were performed immediately after the culture dishes were removed from the incubator.

