## Supplementary Figures for "Ezh2 restrains macrophage inflammatory responses, and is critical for neutrophil migration in response to pulmonary infection"

Figure S1: WT = EZH2 fl/fl KO = LysM x EZH2 fl/fl

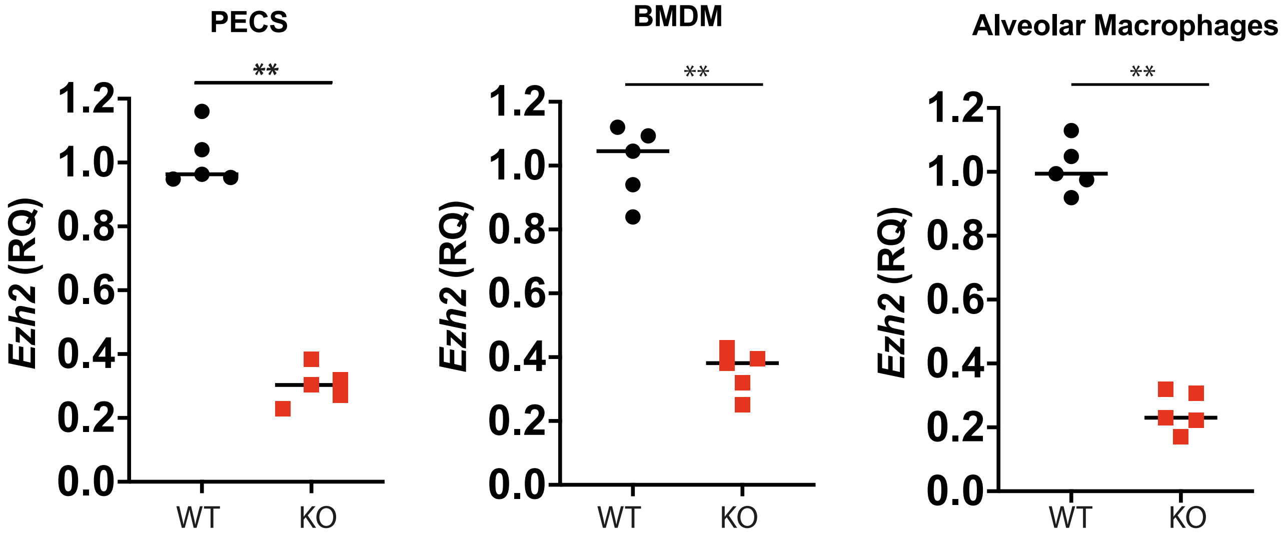

Figure S2

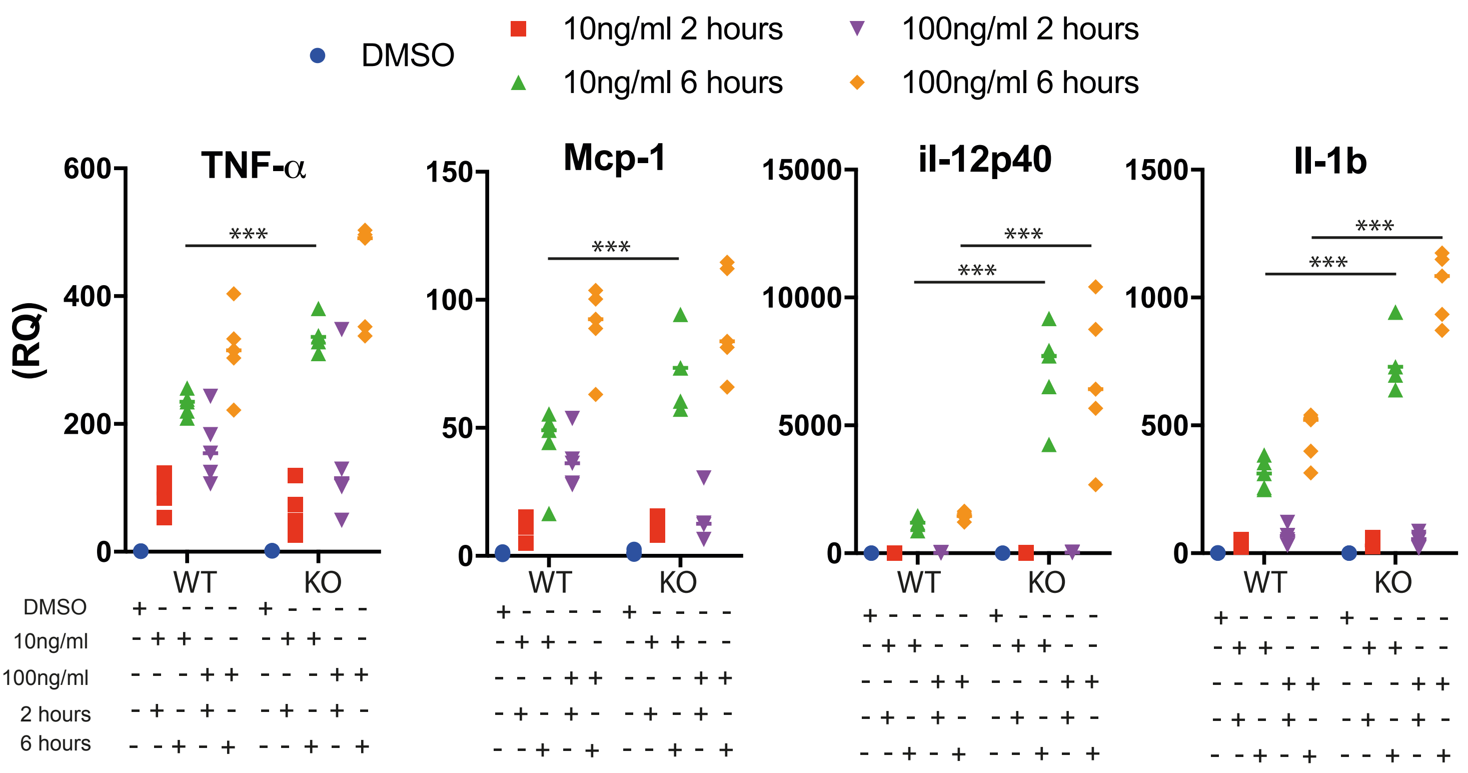
DMSO
10ng/ml 2 hours
10ng/ml 6 hours
100ng/ml 2 hours
100ng/ml 6 hours
TNF-α
Mcp-1
il-12p40
Il-1b
(RQ)
***
WT
KO
DMSO
10ng/ml
100ng/ml
2 hours
6 hours

Figure S3

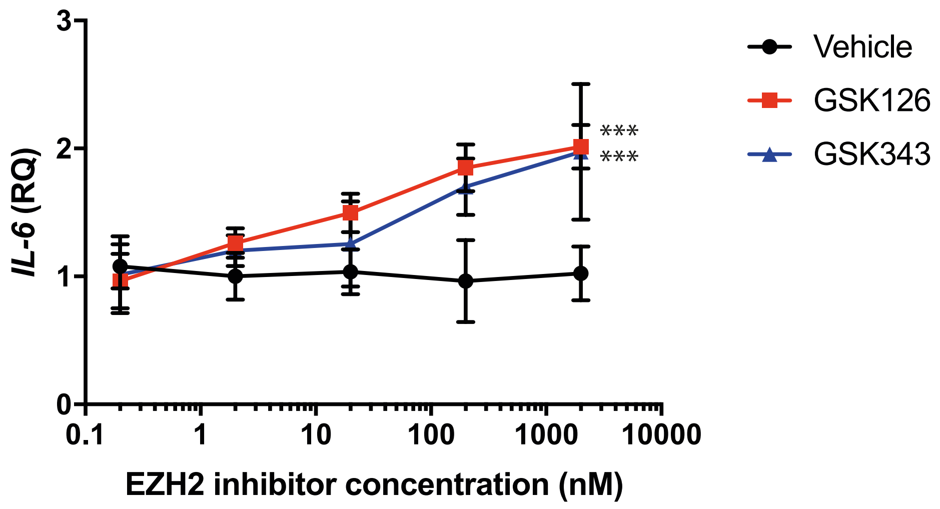

IL-6 (RQ)
3
2
1
0
0.1
1
10
100
1000
10000
EZH2 inhibitor concentration (nM)
***
***
Vehicle
GSK126
GSK343

### Figure S4

| | | | |
|---|---|---|---|
| C1s | Csf2 | Il23a | Nos2 |
| C4a | Csf3 | Il23r | Nox1 |
| C7 | Cxcl1 | Il3 | Nr3c1 |
| Ccl19 | Cxcl10 | Il4 | Oas1a |
| Ccl21a | Cxcl2 | Il5 | Oas2 |
| Ccl8 | Cxcl3 | Il6 | Oasl1 |
| Chi3l3 | Cxcl5 | Il6ra | Pdgfa |
| Defa-rs1 | Cxcl9 | Il7 | Pik3c2g |
| Ifit3 | Cxcr1 | Il9 | Pla2g4a |
| Ifna1 | Cxcr2 | Irf1 | Plcb1 |
| Il1a | Cxcr4 | Irf3 | Ppp1r12b |
| Il22 | Cysltr1 | Irf5 | Prkca |
| Rhoa | Cysltr2 | Irf7 | Prkcb |
| Ager | Daxx | Itgb2 | Ptger1 |
| Alox12 | Ddit3 | Jun | Ptger2 |
| Alox15 | Elk1 | Keap1 | Ptger3 |
| Alox5 | Fasl | Kng1 | Ptger4 |
| Areg | Flt1 | Limk1 | Ptgfr |
| Arg1 | Fos | Lta | Ptgir |
| Atf2 | Fxyd2 | Ltb | Ptgs1 |
| Bcl2l1 | Gnaq | Ltb4r1 | Ptgs2 |
| Bcl6 | Gnas | Ltb4r2 | Ptk2 |
| Birc2 | Gnb1 | Ly96 | Rac1 |
| C1qa | Gngt1 | Maff | Raf1 |
| C1qb | Gpr44 | Mafg | Rapgef2 |
| C1ra | Grb2 | Mafk | Rela |
| C2 | H2-Ea-ps | Map2k1 | Relb |
| C3 | H2-Eb1 | Map2k4 | Retnla |
| C3ar1 | Hc | Map2k6 | Ripk1 |
| C6 | Hdac4 | Map3k1 | Ripk2 |
| C8a | Hif1a | Map3k5 | Rock2 |
| C8b | Hmgb1 | Map3k7 | Rps6ka5 |
| C9 | Hmgb2 | Map3k9 | Shc1 |
| Ccl11 | Hmgn1 | Mapk1 | Smad7 |
| Ccl17 | Hras1 | Mapk14 | Stat1 |
| Ccl2 | Hsh2d | Mapk3 | Stat2 |
| Ccl20 | Hspb1 | Mapk8 | Stat3 |
| Ccl22 | Hspb2 | Mapkapk2 | Tbxa2r |
| Ccl24 | Ifi27l2a | Mapkapk5 | Tcf4 |
| Ccl3 | Ifi44 | Masp1 | Tgfb1 |
| Ccl4 | Ifit1 | Masp2 | Tgfb2 |
| Ccl5 | Ifit2 | Max | Tgfb3 |
| Ccl7 | Ifnb1 | Mbl2 | Tgfbr1 |
| Ccr1 | Ifng | Mef2a | Tlr1 |
| Ccr2 | Iigp1 | Mef2b | Tlr2 |
| Ccr3 | Il10 | Mef2c_Mm | Tlr3 |
| Ccr4 | Il10rb | Mef2d | Tlr4 |
| Ccr7 | Il11 | Mknk1 | Tlr5 |
| Cd163 | Il12a | Mmp3 | Tlr6 |
| Cd4 | Il12b | Mmp9 | Tlr7 |
| Cd40 | Il13 | Mrc1 | Tlr8 |
| Cd40lg | Il15 | Mx1 | Tlr9 |
| Cd55 | Il17a | Mx2 | Tnf |
| Cd86 | Il18 | Myc | Tnfaip3 |
| Cdc42 | Il18rap | Myd88 | Tnfsf14 |
| Cebpb | Il1b | Myl2 | Tollip |
| Cfb | Il1r1 | Nfatc3 | Tradd |
| Cfd | Il1rap | Nfe2l2 | Traf2 |
| Cfl1 | Il1rn | Nfkb1 | Trem2 |
| Creb1 | Il2 | Nlrp3 | Tslp |
| Crp | Il21 | Nod1 | Twist2 |
| Csf1 | Il22ra2 | Nod2 | Tyrobp |

Figure S5a

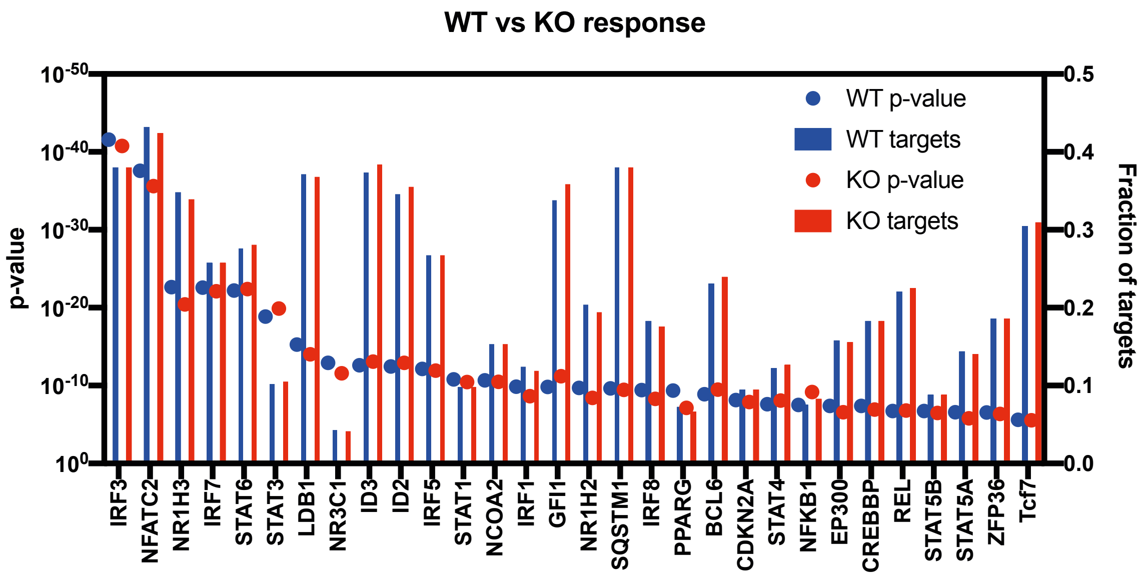

WT vs KO response
p-value
Fraction of targets
WT p-value
WT targets
KO p-value
KO targets
$10^{-50}$
$10^{-40}$
$10^{-30}$
$10^{-20}$
$10^{-10}$
$10^{0}$
0.5
0.4
0.3
0.2
0.1
0.0
IRF3
NFATC2
NR1H3
IRF7
STAT6
STAT3
LDB1
NR3C1
ID3
ID2
IRF5
STAT1
NCOA2
IRF1
GFI1
NR1H2
SQSTM1
IRF8
PPARG
BCL6
CDKN2A
STAT4
NFKB1
EP300
CREBBP
REL
STAT5B
STAT5A
ZFP36
Tcf7

Figure S5b

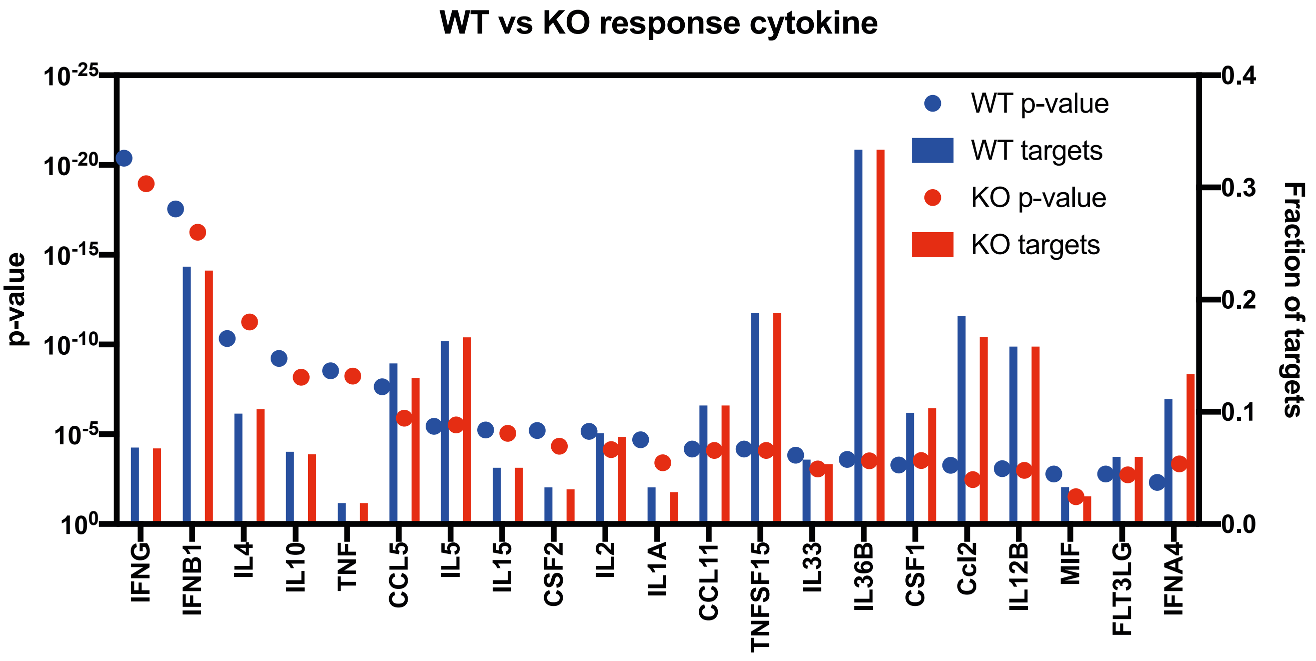
WT vs KO response cytokine
p-value
Fraction of targets
$10^{-25}$
$10^{-20}$
$10^{-15}$
$10^{-10}$
$10^{-5}$
$10^{0}$
0.4
0.3
0.2
0.1
0.0
WT p-value
WT targets
KO p-value
KO targets
IFNG
IFNB1
IL4
IL10
TNF
CCL5
IL5
IL15
CSF2
IL2
IL1A
CCL11
TNFSF15
IL33
IL36B
CSF1
Ccl2
IL12B
MIF
FLT3LG
IFNA4

Figure S5C

#### Basal regulators

| Upstream... | Expr Log ... | Molecule ... | Predicted ... | Activation ... | Flags | p-val... | Target Mo... | Mechanist... |
|---|---|---|---|---|---|---|---|---|
| PPARG | ↑0.540 | ligand-depende... | | 0.803 | | 5.41E-03 | ↑ABCG1, ....all 19 | |
| Tcf7 | ↓-1.039 | transcription reg... | | -1.704 | | 6.81E-03 | ↑CCR2,↓.....all 31 | |
| ID3 | ↓-0.620 | transcription reg... | | -0.853 | | 7.80E-03 | ↓ADM, ↑......all 37 | |
| Meis1 | ↓-0.758 | transcription reg... | | | | 2.34E-02 | ↑HIF1A,↓......all 2 | |

#### LPS regulators

| Upstream R... | Expr Log Ratio | Molecule Type | Predicted Ac... | Activation z-... | p-value ... | Target Mole... |
|---|---|---|---|---|---|---|
| IRF7 | ↑2.103 | transcription regula... | Activated | 5.562 | 1.41E-18 | ↑CCL5,↑C... ...all 33 |
| BCL3 | ↑0.557 | transcription regula... | Inhibited | -2.597 | 4.85E-06 | ↑CD69, ↓... ...all 12 |
| CEBPE | ↓-0.911 | transcription regula... | | 1.519 | 1.98E-04 | ↑Ccl7, ↑CD14...all 8 |
| SPIB | ↑3.883 | transcription regula... | | 1.000 | 6.73E-04 | ↓APOE, ↑... ...all 16 |
| PPARA | ↑1.175 | ligand-dependent ... | | -0.933 | 2.20E-03 | ↓ABCA1, ↑... ...all 6 |
| NFKBIZ | ↑2.117 | transcription regula... | | 1.953 | 9.14E-03 | ↑Ccl2, ↑CSF2, ...all 4 |
| NFKBIA | ↑1.519 | transcription regula... | | | 2.41E-02 | ↑CCL5, ↑C... ...all 6 |
| IRF1 | ↑1.198 | transcription regula... | | 1.981 | 2.43E-02 | ↑BCL2, ↑CC... ...all 8 |
| CBFA2T3 | ↓-0.563 | transcription regula... | | | 2.49E-02 | ↓CD4, ↑ITG... ...all 3 |

### Figure S5D

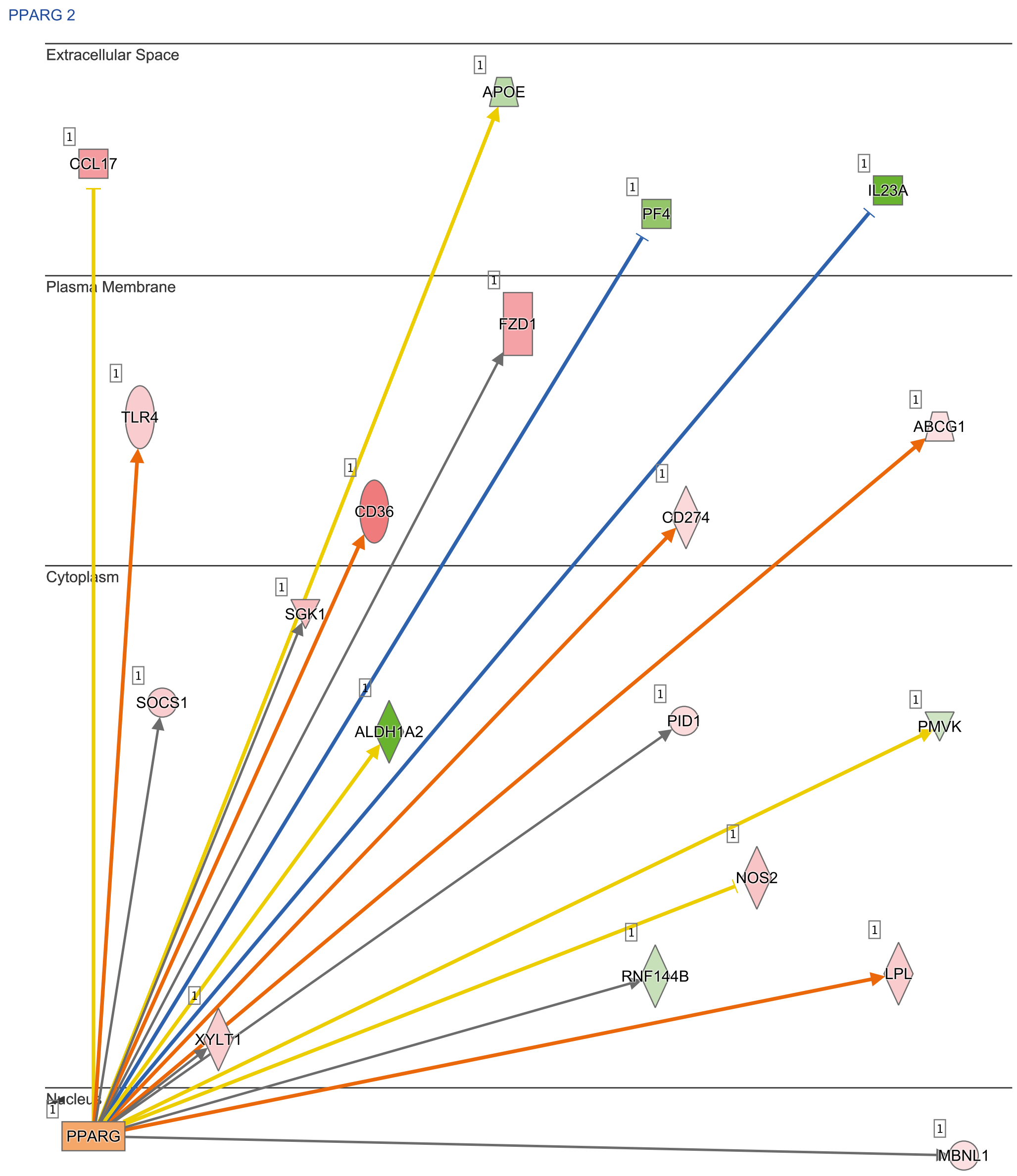


© 2000-2020 QIAGEN. All rights reserved.

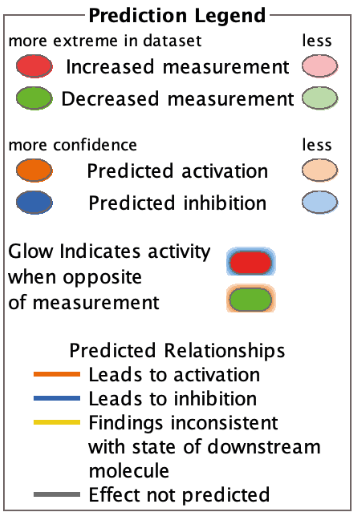

Figure S5E

BCL3,IRF1,IRF7,NFKBIA,NFKBIZ,PPARA,SPIB 1

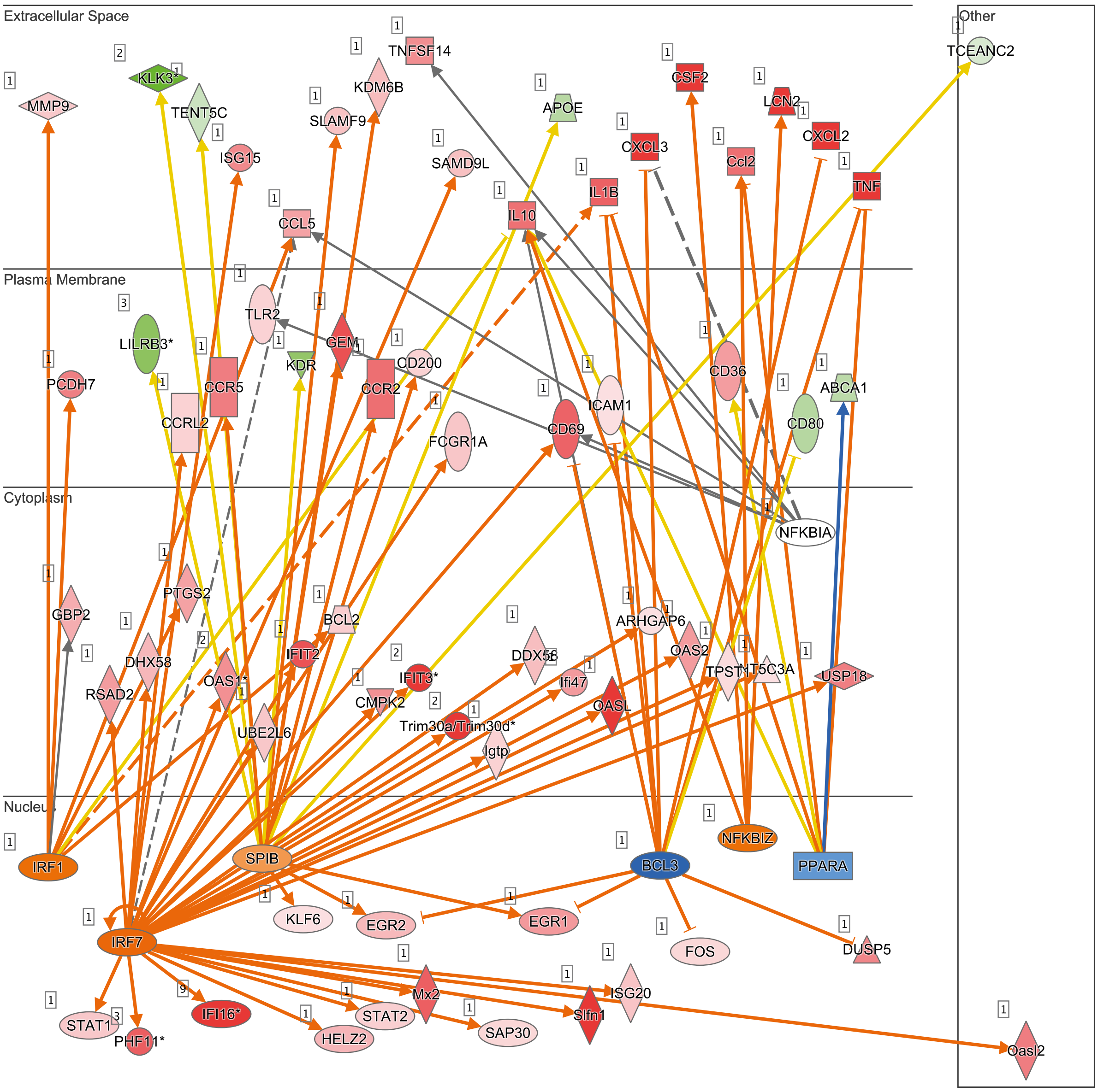


© 2000-2020 QIAGEN. All rights reserved.

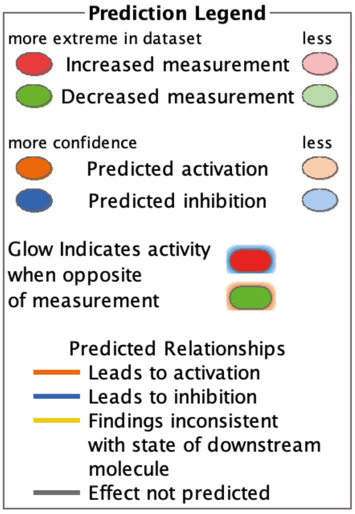

Figure S6

A

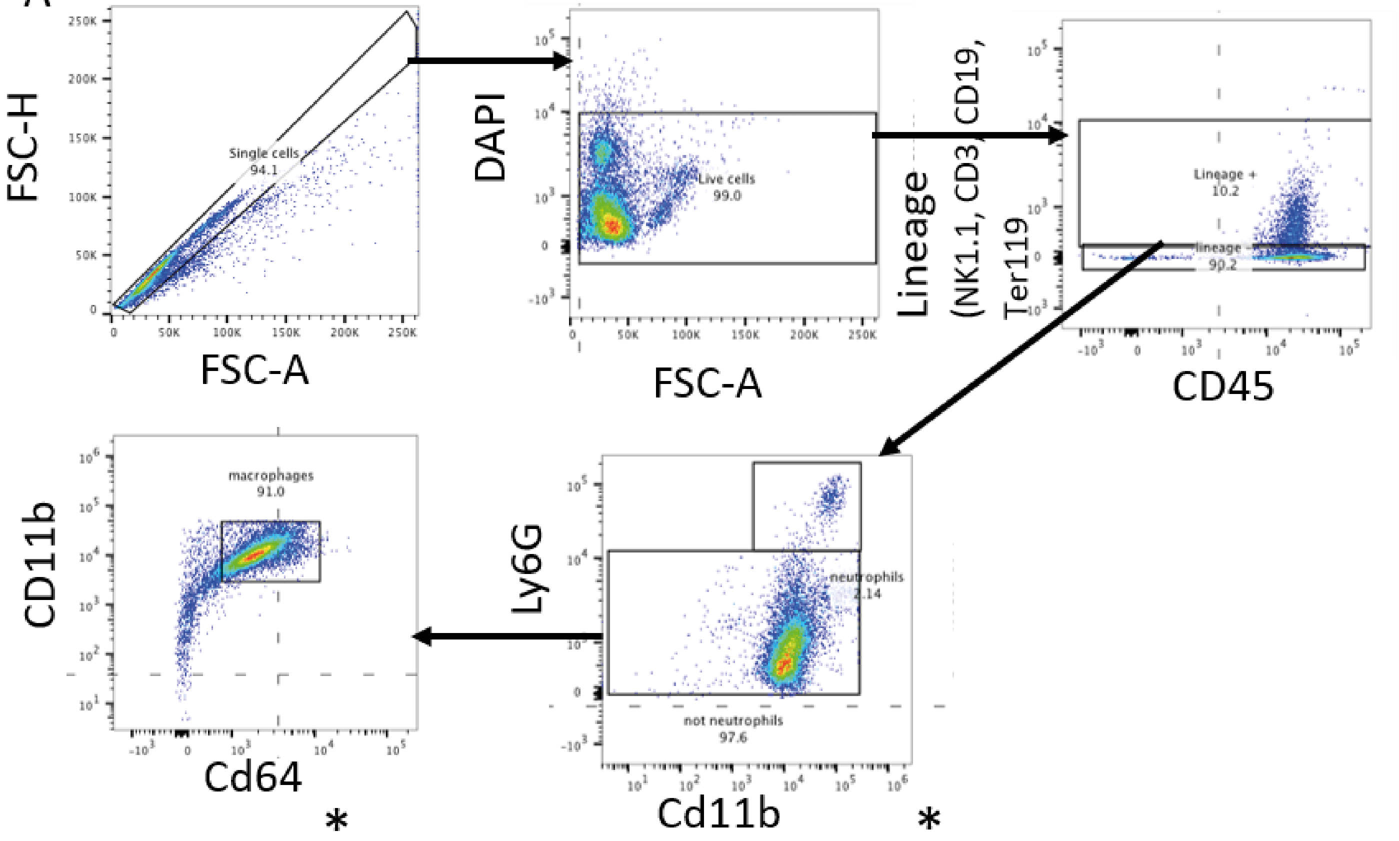

FSC-H
FSC-A
Single cells
94.1
DAPI
Live cells
99.0
Lineage (NK1.1, CD3, CD19, Ter119
CD45
Lineage +
10.2
lineage -
90.2
Ly6G
Cd11b
neutrophils
2.14
not neutrophils
97.6
CD11b
Cd64
macrophages
91.0
*
*


B

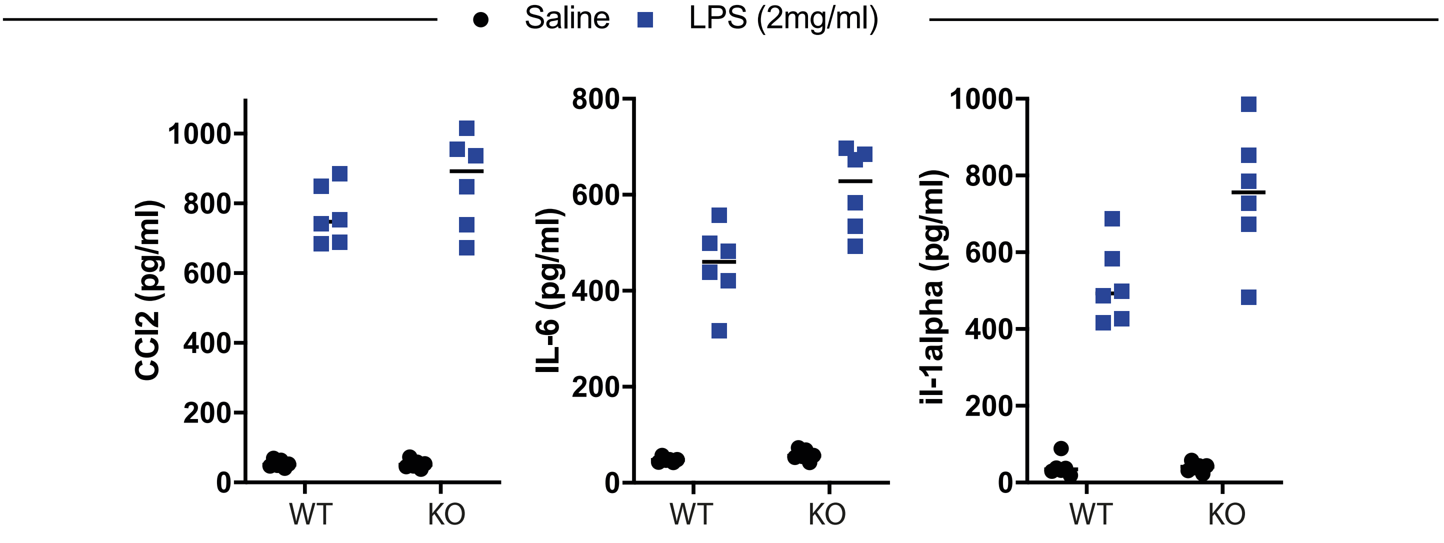

Saline
LPS (2mg/ml)
Ccl2 (pg/ml)
IL-6 (pg/ml)
il-1alpha (pg/ml)
WT
KO

Figure S7

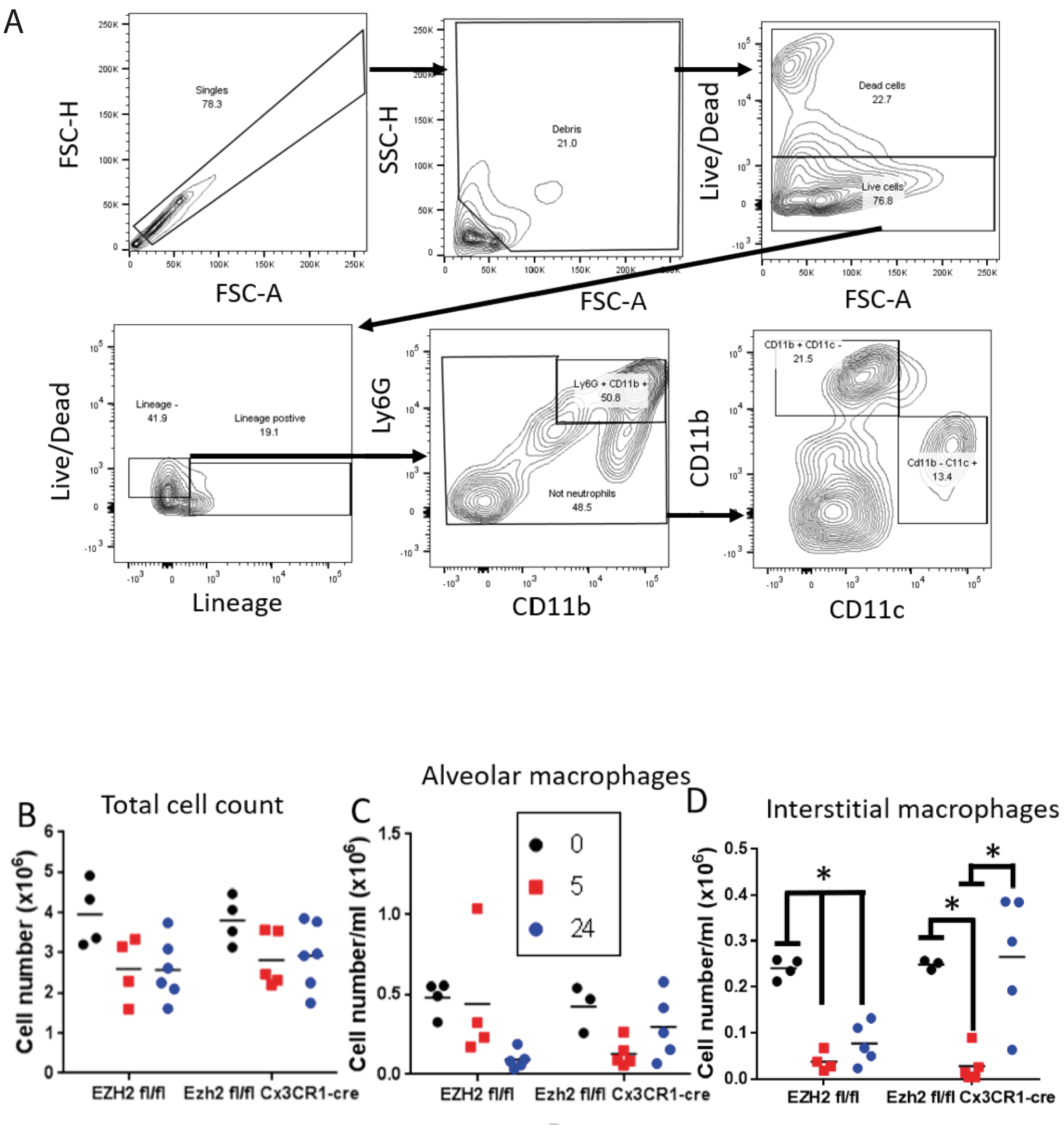

A
FSC-H
FSC-A
Singles
78.3
SSC-H
Debris
21.0
Live/Dead
Dead cells
22.7
Live cells
76.8
Lineage -
41.9
Lineage postive
19.1
Lineage
Ly6G
Ly6G + CD11b +
50.8
Not neutrophils
48.5
CD11b
CD11b + CD11c -
21.5
Cd11b - C11c +
13.4
CD11c
B
Total cell count
Cell number (x10⁶)
EZH2 fl/fl
Ezh2 fl/fl Cx3CR1-cre
C
Alveolar macrophages
Cell number/ml (x10⁶)
0
5
24
D
Interstitial macrophages
*

Figure S8

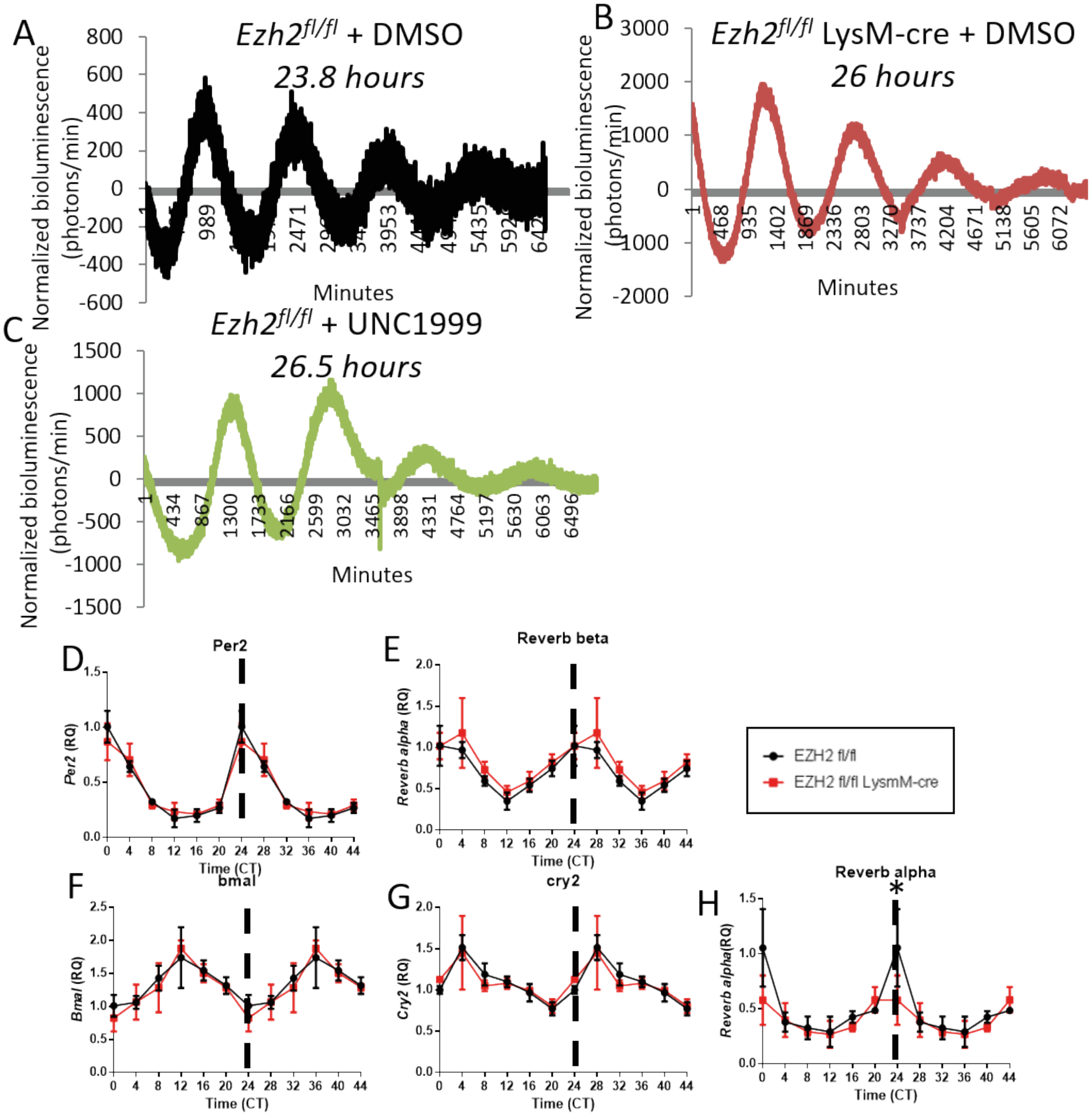
A
Ezh2fl/fl + DMSO
23.8 hours
Normalized bioluminescence (photons/min)
Minutes
B
Ezh2fl/fl LysM-cre + DMSO
26 hours
C
Ezh2fl/fl + UNC1999
26.5 hours
D
Per2
Time (CT)
E
Reverb beta
Reverb alpha (RQ)
EZH2 fl/fl
EZH2 fl/fl LysmM-cre
F
bmal
Bmal (RQ)
G
cry2
Cry2 (RQ)
H
Reverb alpha
*
