## Supplementary Figure legends for "Ezh2 restrains macrophage inflammatory responses, and is critical for neutrophil migration in response to pulmonary infection"

Supplementary legends

FigS1

Transcript concentration for Ezh2, from peritoneal exudate cells (PECs), Bone marrow derived macrophages (BMDM), and alveolar macrophages. (n=5, median plotted, Mann-Whitney *U*-test **p<0.01, ***p<0.001)

Fig S2

Isolated macrophages from bone marrow and treated with different doses of LPS for 2 or 6 hours shows that these cells are highly sensitive to time and concentration of treatment. Ex-vivo BMDMs from LysM.EZH2^-/-^ macrophages and littermate controls were treated with LPS for 2 or 6 hours at a dose of 10 or 100ng/ml and the cytokine response assessed by RNA quantification. (n= 5, median plotted, 2-way ANOVA, post-hoc Tukey’s multi-comparison test. *** p<0.001).

Fig S3

EZH2 inhibitor dose response curve. Transcript abundance of Il-6 in ex-vivo bone marrow derived macrophages treated for 2hr with LPS (100ng/ml) following 48 hours incubation with one of the EZH2 inhibitors BSK343, GSK126, or DMSO control. IL-6 mRNA was quantified by qRT-PCR relative to Gaphd, and normalised to DMSO vehicle treated bone marrow derived macrophages. (n= 3, 2-way ANOVA, post-hoc Tukey’s multi-comparison test. *** p<0.001).

Fig S4

Gene list for the Mouse inflammation panel V2 Nanostring^TM^

Fig S5A

Upstream regulators responsible for differential gene expression following LPS treatment were identified for EZH2^f/f^ (blue) and LysM-cre.EZH2^fl/fl^ (red) macrophage cells using Ingenuity Pathway Analysis (Qiagen). Genes were considered to be differentially expressed if log2 fold change had a magnitude greater than 1 and the adjusted p-value of differential expression was <1x10^-5^. Plot shows the p-value of enrichment (points) and proportion of targets which were differentially expressed (bars) in response to LPS treatment. The top 30 upstream regulators belonging to either the ‘transcriptional regulator’ or ‘ligand-binding nuclear receptor’ categories are shown (30/30 overlap).

S5B

Plot showing the p-value of enrichment (points) and proportion of targets which were differentially expressed (bars) in response to LPS treatment for differentially expressed genes as in S5A. The top 20 upstream regulators belonging to the ‘cytokine’ category are shown (19/20 overlap between WT and KO conditions).

S5C

Upstream regulators responsible for differential gene expression between EZH2^fl/fl^ and LysM-cre.EZH2^fl/fl^ macrophages under basal conditions (top) and LPS-treated conditions (bottom) were identified using Ingenuity Pathway Analysis (Qiagen). Genes were considered to be differentially expressed if adjusted p-value was <0.05. Upstream regulators belonging to either the ‘transcriptional regulator’ or ‘ligand-binding nuclear receptor’ categories are shown where the magnitude of expression log ratio was greater than 0.5.

S5D

Network diagram showing an example upstream transcriptional regulator (PPARg) differentially enriched in LysM-cre.EZH2^f/f^ macrophages under basal conditions, with differentially expressed downstream targets.

S5E

Network diagram showing upstream transcriptional regulators differentially enriched in LysM-cre.EZH2^f/f^ macrophages under LPS-treated conditions, with differentially expressed downstream targets.

Fig S6

A) Flow cytometry gating strategy for quantification of pulmonary infiltrate quantified in figure 4B

B) Pulmonary in-vivo inflammatory responses in CX3CR1-cre EZH2^fl/fl^ (KO) and EZH2^fl/fl^ (WT) mice. Mice were subject to nebulized LPS, and after 4 hours killed. The bronchoalveolar lavage (BAL) was analysed and the cytokine values are shown(n=6, median plotted, 2-way ANOVA, post-hoc Tukey’s multi-comparison test. P value ns)

Fig S7

Conditional deletion of EZH2 in macrophages alters the re-population of interstitial macrophages after aerosolised LPS. (A) flow cytometry identification marks used to identify immune cells. (B) total cell counts, (C) Cd11c +Cd11b – alveolar macrophages and (D) cd11c- Cd11b+ interstitial macrophages found in lung digest after BAL, 0, 5, and 24 hours after a 20 minute exposure to 2mg/ml aerosolised LPS challenge in EZH2 ^fl/fl^.CX3CR1 and floxed control mice. (n= 5, median plotted, 2-way ANOVA, post-hoc Tukey’s multi-comparison test. * p<0.05).

Fig S8

The loss of EZH2 alters the period of the circadian oscillator. (A) an example of the bioluminescence recordings from PECs cultures from EZH2^fl/fl^ mice (black). PECs derived from EZH2 ^fl/fl^.LysMcre mice (red) and (C) PECs cultured from EZH2^fl/fl^ mice treated with 2uM UNC1999 for 48 hours prior to synchronisation (green). All mice were on a PER2::LUC background. The luciferase activity is measured in live cells using recoding medium containing D-Luciferin. Rhythmicity was determined using the JTK package, and the calculated circadian period is indicated.

The transcripts of the core circadian clock genes were analysed in temperature synchronised bone marrow derived macrophages harvested from EZH2 ^fl/fl^.LysMcre mice and floxed controls. The genes measured were (D) Per2, (E) ReverbB, (F) Bmal1, (G) Cry2, and (H) Reverba. The genes were measured at 4 hourly intervals. Profiles are double plotted, mRNA was quantified relative to GAPDH and is normalised to values at CT0 in control cells. The data is presented as the mean (n=3 per time point, *=p value <0.05 vs control cells)
